# A Comprehensive Collection of Pain and Opioid Use Disorder Compounds for High-Throughput Screening and Artificial Intelligence-Driven Drug Discovery

**DOI:** 10.1101/2024.04.21.590471

**Authors:** Xin Hu, Paul Shinn, Zina Itkin, Lin Ye, Ya-Qin Zhang, Min Shen, Stephanie Ford-Scheimer, Matthew D. Hall

## Abstract

As part of the NIH Helping to End Addiction Long-term (HEAL) Initiative, the National Center for Advancing Translational Sciences (NCATS) is dedicated to the development of new pharmacological tools and investigational drugs for managing and treating pain, as well as the prevention and treatment of opioid misuse and addiction. In line with these objectives, we created a comprehensive, annotated small molecule library including drugs, probes, and tool compounds that act on published pain and addiction-relevant targets. Nearly 3,000 small molecules associated with approximately 200 known and hypothesized HEAL targets have been assembled, curated, and annotated in one collection. Physical samples of the library compounds have been acquired and plated in 1536-well format, enabling rapid and efficient high throughput screen (HTS) against a wide range of assays. The creation of the HEAL Targets and Compounds Library, coupled with an integrated computational platform for AI-driven machine learning (ML), structural modeling, and virtual screening (VS), provides a valuable source for strategic drug repurposing, innovative profiling, and hypothesis testing of novel targets related to pain and opioid use disorder (OUD). The library is available to investigators for screening pain and OUD-relevant phenotypes.

---

The opioid crisis highlights the urgent need for novel non-addictive pain medications, as well as improved treatments for opioid misuse and addiction ^1, 2^. Advances in our understanding of the molecular and neural circuit physiology of pain and opioid use disorder (OUD), have unveiled many putative approaches for therapeutic intervention, such as pharmacological agents directed at novel targets to test new therapeutic hypotheses, the development of new drugs aimed at these targets, and the establishment of new testing systems with the potential to offer more accurate predictions of human clinical response compared to traditionally used models ^3, 4^.

Significant challenges remain for the creation of new therapeutics ^5, 6^, largely due to the absence of drug-like pharmacological probes for novel targets, hindering the exploration of new therapeutic hypotheses. Additionally, the preclinical models for pain and addiction need to be refined to predict efficacy more reliably in humans, and the market attractiveness for addiction and overdose treatments is currently limited ^7, 8^.

The Helping to End Addiction Long-term (HEAL) Initiative, launched in April 2018 as a collaborative effort across the National Institutes of Health (NIH), is dedicated to advancing national priorities in combating the opioid crisis through scientific endeavors ^9–11^. As part of the HEAL Initiative, the National Center for Advancing Translational Sciences (NCATS) is committed to accelerating the process of discovery and demonstration of new treatments for opioid misuse, addiction, and pain. Our collaborative program actively engages in the development of novel chemistry, screening, and testing methodologies to discover new pharmacological probes and investigational drugs tailored for addressing the challenges of pain and OUD.

Pharmacogenomics can be a valuable research approach for developing and evaluating cell-based models of pathologies such as those associated with the HEAL Initiative. This approach requires the availability of a library of small molecules with annotated mechanisms of action whose activity in biological assays can be related back to target biology. Ideally, such annotated libraries should contain a variety of molecules with redundant mechanisms of action to enhance confidence in their activity, and the target space covered by molecules present in the library is typically sourced from the existing literature. Beyond providing insight into biological mechanisms, screening with annotated libraries also offers the opportunity to identify potential drug repurposing candidates for the treatment of specific disorders. A range of annotated libraries have been reported, each designed with a particular focus. Some libraries are aiming at approved therapeutics (for example, the NCATS Pharmaceutical Collection, NPC ^12^) or general coverage of clinical space (such as the Broad Drug Re-purposing Hub ^13^). Annotated libraries have also been designed around specific disease biology target space, as seen in the case of the NCATS MIPE library, which contains molecules with defined mechanisms of action associated with oncology indications ^14^.

As of now, there is no annotated library specifically designed to address the pain and addiction biological space. To fill this gap, we have created a comprehensive annotated HEAL library by curating known and potential targets related to pain and OUD, as well as assembling known drugs, chemical probes, and tool compounds associated with each of these targets. The creation of the HEAL Targets and Compounds Library incorporates valuable insights gained from high-throughput screening and chemical library design at NCATS. This best-in-class repurposing library enables rapid testing of drug candidate compounds with established safety profiles in the context of pain and addiction, as well as the identification of targets and pharmacology associated with mediating pain and addiction phenotypes. The collection can be made available in pre-spotted plates for investigators with laboratory assays able to test against novel biology in 384-well plates. Moreover, the HTS data generated at NCATS with this library will be openly shared with the scientific community, providing a valuable resource for researchers.

Here, we provide detailed information on the creation and composition of the HEAL library. A case study on several HEAL targets using quantitative HTS (qHTS) and virtual screening (VS) for probe and drug discovery as part of the collaborative program at NCATS is summarized. This case study illustrates the practical application of the HEAL targets and compounds as an integrated platform for AI-driven machine learning, *in silico* structural modeling, and large-scale database virtual screening for therapeutics development against pain and OUD.

## Creating the HEAL Library target list

For the purposes of the HEAL Library design, potential HEAL targets could be broadly defined to include any targets that are involved in opioid and pain signaling pathways and implicated as a regulator of pain and inflammation. A survey of validated protein targets for addiction and pain in DrugBank ^15^ and ChEMBL ^16^ showed that only about 30-40 small molecule drug targets are associated with pain and addiction. These classical analgesic targets that provide clinically validated efficacy are mostly nociceptors and neurological targets, such as opioid receptors, dopamine receptors, serotonin receptors, adrenoceptors, cannabinoid receptors, and ligand-gated and voltage-gated ion channels. To tackle the problems and side effects associated with these traditional drugs and targets, novel approaches were developed, such as the opioid receptors that block pain but don’t affect breathing or reward signaling, or alternative opioid receptors that are not located in the reward center of the brain ^5^. On the other hand, a large number of non-opioid pain targets have been identified for the development of novel analgesics. These novel targets have been reviewed recently ^17, 18^, including P2X3 and P2X7 purinoceptors, NaV1.7 and NaV1.8 sodium-channels, orphan receptors GPR18/GPR35/GPR55, enzyme Triointsub receptor kinase A (TrkA), sepiapterin reductase (SPR), and soluble epoxide hydrolase (sEH).

In addition to assembling these classical pain and OUD-related targets in the HEAL Library, we also included a broader collection of novel and hypothesized targets that have been investigated for pain and OUD treatment in the literature. For example, PIEZO2 is a stretch-gated cell membrane ion channel expressed in sensory neurons, nociceptors, and mechanosensitive epithelial cells. Targeting PIEZO2 provides novel ways to treat tactile pain and chronic itch while preserving normal pain and temperature sensation ^19^. The galanin 1 receptor (GAL1R) forms heteromers with µ-opioid receptor (MOR) in the ventral tegmental area that modulates dopamine release. GAL1R ligands block MOR-mediated signaling by allosteric modulations within the MOR-GAL1R heterodimer. Targeting MOR-GAL1R heterodimers is an appealing strategy for novel drug design to treat OUD while sparing the analgesic effect of opioids ^20^. Some novel targets that are currently under investigation for high throughput screening for probe and drug development at NCATS are listed in **Table 1**. A list of all targets, including novel and hypothesized targets used for the creation of the HEAL Library, is provided (**Supplementary Table S1**). The details of library composition are discussed in the following section.

**Table 1.** HEAL targets for pain and OUD.

| HEAL Targets | Background | MOA | Status of Research |
| --- | --- | --- | --- |
| GAL1R | Galanin receptor 1 (GAL1R) forms heteromers with the μ-opioid receptor (MOR) in the ventral tegmental area (VTA) that modulate dopamine release. GAL1R ligands block MOR-mediated signaling by allosteric modulations within the MOR-GAL1R heteromers. | Targeting MOR-GAL1R heteromers to treat OUD while sparing the analgesic effect of opioids. | Structure of GAL1R bound with galanin peptide available. No specific small molecules against GAL1R reported. |
| GPR151 | GPR151 is an orphan GPCR almost exclusively in the medial habenula (mHb), which is co-localized with $\mu$ opioid receptors. GPR151 modulates opioid receptor signaling in mHb. | Modulating opioid signaling in mHb by targeting GPR151 signaling as a novel treatment for opioid dependence in humans. | No ligand and small molecule agonists or antagonists reported. |
| RXFP3 | Relaxin-3 is a neuropeptide that is highly expressed in the brain with important roles in metabolism, arousal, learning and memory. Its cognate receptor relaxin-3 receptor (RXFP3) is involved in multiple disorders including reward and emotional processing, stress responses, cognition and drug addiction. | RXFP3 agonists are a potential treatment for pain, anxiety, depression, and OUD. | A series of small molecule agonists reported. Probe compounds for family targets RXFP1 and RXFP2 identified. |
| PAC1R | The pituitary adenylate cyclase-activating polypeptide type I receptor (PAC1R) is widely distributed in the central neural system and peripheral organs. Abnormal activation of the receptor mediates trigeminovascular activation and sensitization is highly related to migraine. | PAC1R antagonists are potential therapeutics for neuronal disorders. | Structure of PAC1R bound with ligand PACUP peptide available. A series of small molecule antagonists reported. |
| PIEZO2 | PIEZO2 is a stretch-gated cell membrane ion channel expressed in sensory neurons, nociceptors, and mechanosensitive epithelial cells. Loss-of-function produces loss of proprioception, gentle touch, and mechanical allodynia. | PIEZO2 agonists provide novel ways to treat tactile pain and chronic itch while preserving normal pain and temperature sensation. | Structure of full-length PIEZO2; no agonist/antagonist reported. A series of small molecule agonists identified for PIEZO1. |
| NPR1 | Atrial natriuretic peptide receptor 1 (NPR1) is a receptor for the brain natriuretic peptide NPPA, which is a critical component required for the excitation of spinal cord *Npr1*-expressing interneurons. | Inhibition of *Npr1* represents a novel strategy for the treatment of acute and chronic itch. | Small molecule antagonists reported but none in development. |
| AC1 | Adenyl cyclase 1 (AC1) is highly expressed in neuronal tissues associated with pain processing and neuronal plasticity. Knockout animals reveal almost a complete loss in pain responsiveness in both inflammatory and neuropathic pain models. | AC1-selective small molecule inhibitors are novel, non-opioid treatment for chronic pain. | Small molecule inhibitors reported but none in development. |
| GCPII | Glutamate carboxypeptidase II (GCPII) is an extracellular enzyme that hydrolyzes NAAG to *N*-acetylaspartate and glutamate. Excess glutamatergic transmission is associated with acute and chronic pain caused by inflammation or brain injury. | GCPII inhibitors are a major unrealized promise for the treatment of pain associated with glutamate excitotoxicity by reducing glutamate levels without interfering with glutamatergic transmission. | Structures and small molecule inhibitors available but none in development. |

## Identification of compounds included in the HEAL Library

As a first step towards creating the HEAL Library, we included 430 unique drugs, including experimental or clinical candidates with in-scope indications in pain that were found in DrugBank and NCATS Inxight Drugs ^21^, along with a few drugs with indications for treating addiction, dependency, and overdose. The targets of these FDA-approved drugs are mostly classical pain targets, such as the opioid and dopamine receptors, cannabinoid receptor, and ion channels. Some of the pain and addiction-relevant drugs that do not have a clear protein target were also included in the HEAL Library. Secondly, new drug candidates in clinical trials associated with all HEAL targets were collected from public resources, including ChEMBL and patent search. **Table 2** shows some examples of HEAL drug candidates in clinical trials. Compounds without disclosed structures (e.g., drugs in Phase 1 clinical trials that have not released the structures of clinical candidates) were annotated in the target library list for future updates (HEAL Library 2.0) once these molecules are commercially available.

**Table 2.**
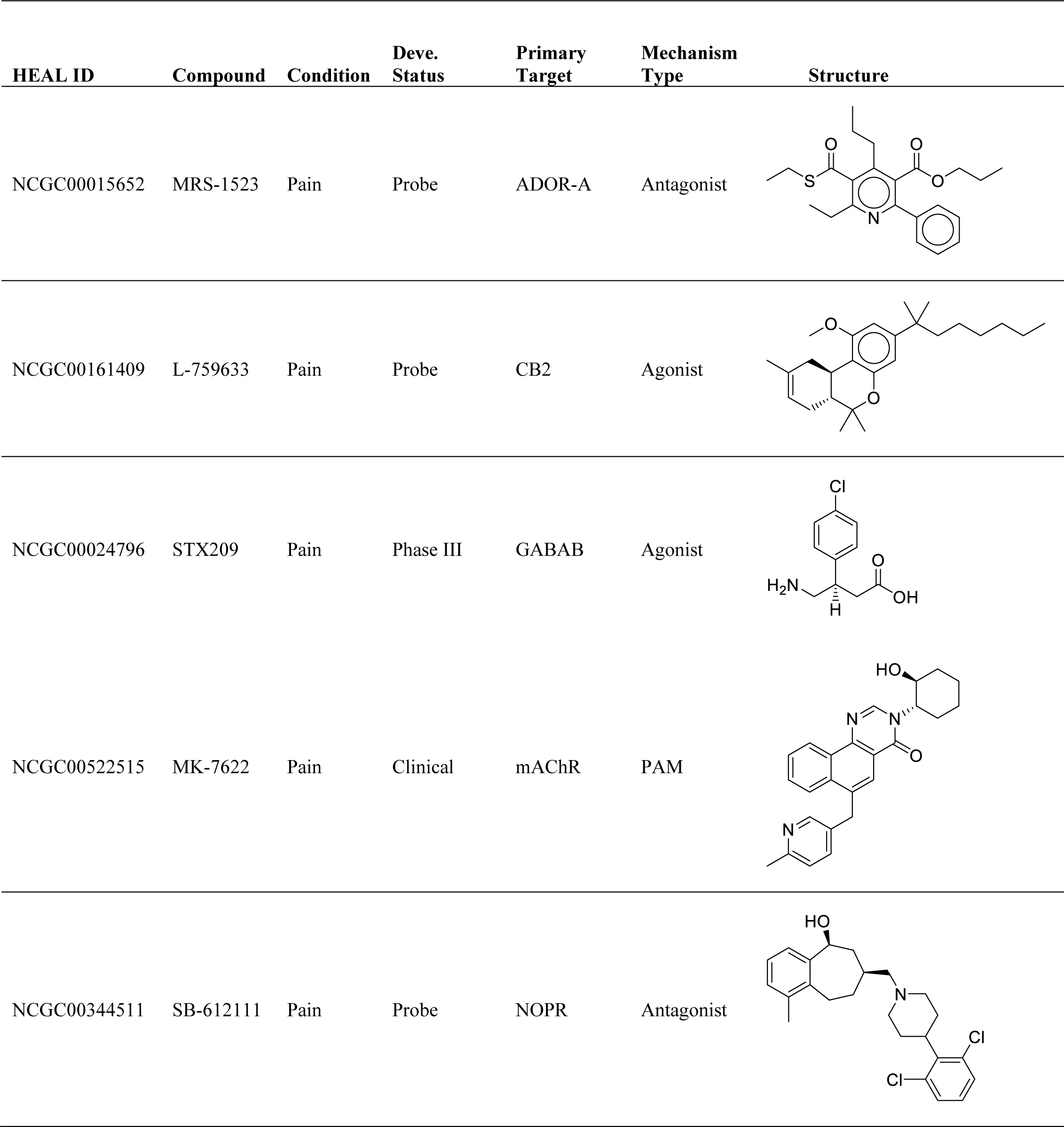
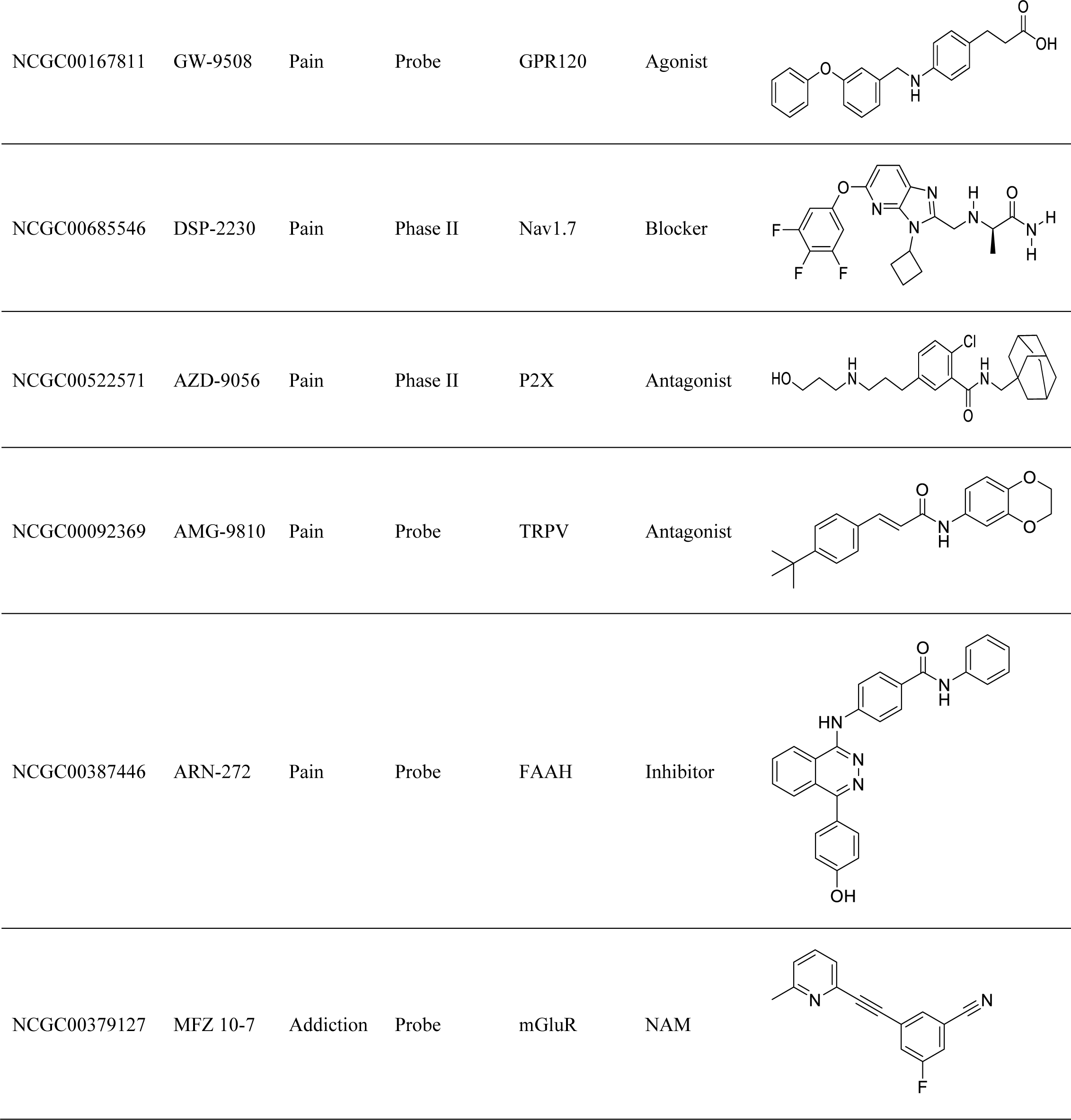
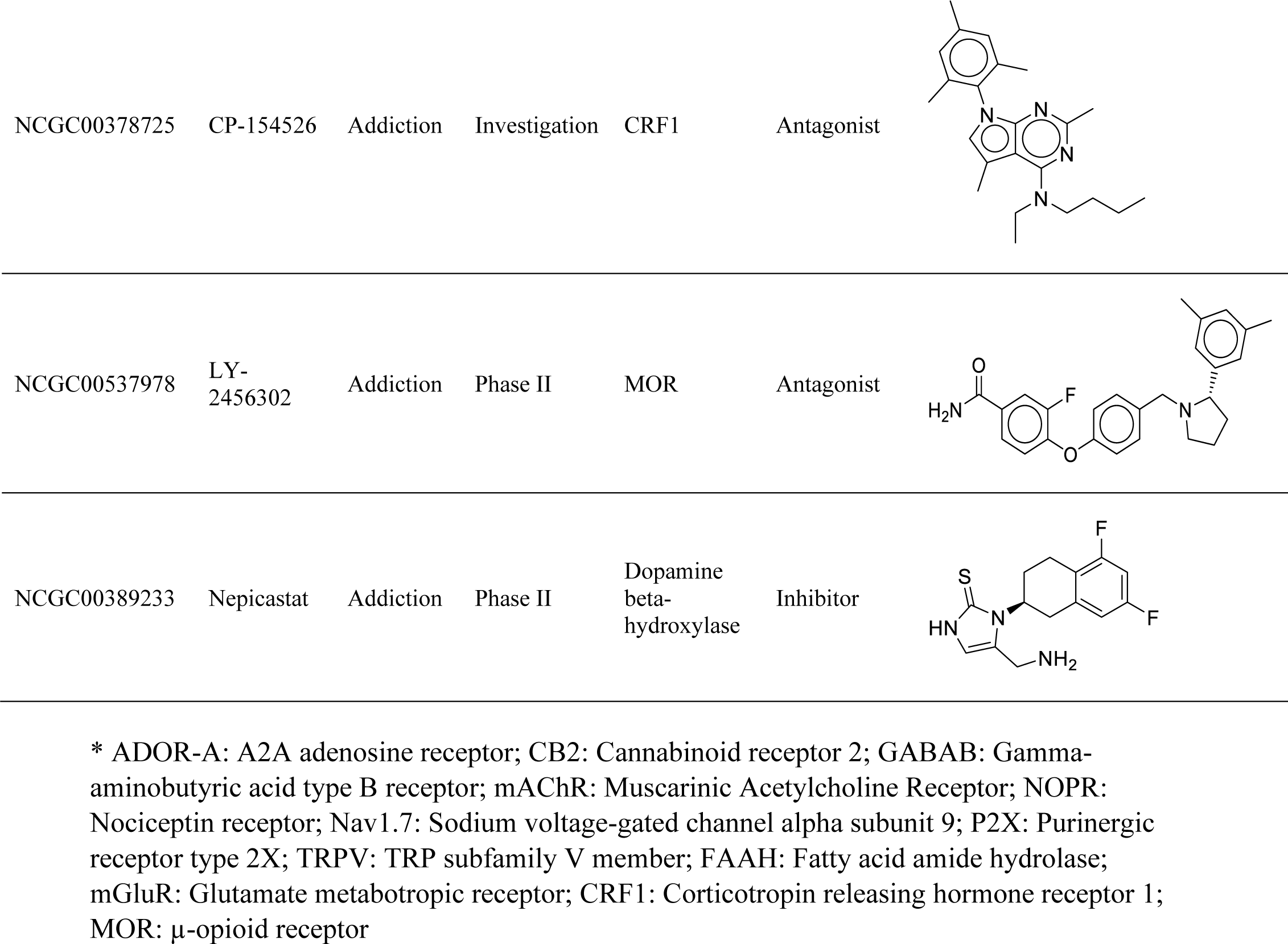
Representatives of HEAL library targets and compounds.

| HEAL ID | Compound | Condition | Deve.<br>Status | Primary<br>Target | Mechanism<br>Type | Structure |
| --- | --- | --- | --- | --- | --- | --- |
| NCGC00015652 | MRS-1523 | Pain | Probe | ADOR-A | Antagonist |  |
| NCGC00161409 | L-759633 | Pain | Probe | CB2 | Agonist |  |
| NCGC00024796 | STX209 | Pain | Phase III | GABAB | Agonist |  |
| NCGC00522515 | MK-7622 | Pain | Clinical | mAChR | PAM |  |
| NCGC00344511 | SB-612111 | Pain | Probe | NOPR | Antagonist |  |
| NCGC00167811 | GW-9508 | Pain | Probe | GPR120 | Agonist |  |
| NCGC00685546 | DSP-2230 | Pain | Phase II | Nav1.7 | Blocker |  |
| NCGC00522571 | AZD-9056 | Pain | Phase II | P2X | Antagonist |  |
| NCGC00092369 | AMG-9810 | Pain | Probe | TRPV | Antagonist |  |
| NCGC00387446 | ARN-272 | Pain | Probe | FAAH | Inhibitor |  |
| NCGC00379127 | MFZ 10-7 | Addiction | Probe | mGluR | NAM |  |
| NCGC00378725 | CP-154526 | Addiction | Investigation | CRF1 | Antagonist |  |
| NCGC00537978 | LY-2456302 | Addiction | Phase II | MOR | Antagonist |  |
| NCGC00389233 | Nepicastat | Addiction | Phase II | Dopamine<br>beta-<br>hydroxylase | Inhibitor |  |

\* ADOR-A: A2A adenosine receptor; CB2: Cannabinoid receptor 2; GABAB: Gamma-aminobutyric acid type B receptor; mAChR: Muscarinic Acetylcholine Receptor; NOPR: Nociceptin receptor; Nav1.7: Sodium voltage-gated channel alpha subunit 9; P2X: Purinergic receptor type 2X; TRPV: TRP subfamily V member; FAAH: Fatty acid amide hydrolase; mGluR: Glutamate metabotropic receptor; CRF1: Corticotropin releasing hormone receptor 1; MOR: μ-opioid receptor

Finally, we searched for small molecule tool compounds acting against targets in the curated target list from various public resources, like PubChem ^22^, BindingDB ^23^, and IUPHAR/BPS ^24^. All pharmacologic mechanisms of action (inhibitors, antagonist, agonist, and modulators) were included in this search, which resulted in an extensive assembly of compounds (over 4,000). To prioritize the list, all compounds were manually annotated and curated.

Structural clustering and molecular properties were evaluated to facilitate the selection of compounds with CNS properties in addition to those implicated in pain/OUD pathways and neurodegenerative or inflammatory diseases. Additionally, we considered the availability of identified small molecules from NCATS in-house libraries and commercial vendors. This process resulted in the identification of approximately 2,400 compounds: 850 were from the approved and clinical drugs list; 490 from the CNS drug-likeness list; and 1,033 from tool and probe molecules acting against targets of interest. Initially, many of them were identified from various disease areas, such as oncology and infectious diseases, reinforcing the value of creating small molecule probes for every protein in the human genome ^25^.

## Annotation of the HEAL Library

The workflow for creating the HEAL Library is illustrated in **Figure 1**. Like other NCATS annotated screening libraries ^12, 26, 27^, all compounds and targets were thoroughly curated, annotated, and deposited into NCATS probeDB, which is managed through a HTS client tool with functionalities for searching, retrieving, updating, and structural analysis. Currently, nearly 3,000 drugs and probe compounds associated with approximately 240 known and hypothesized HEAL-relevant targets have been collected and plated in the HEAL Library (version 1.0). **Figure 2** shows the components of HEAL targets and compounds whose physical samples were plated for HTS (2,800 compounds). Forty-nine percent (49%, 1,366) of the plated library compounds were from the GPCR targets, and 25% (703) were from the ion channels. The least number of compounds was found with transporters (1%) and catalytic receptors (2%), while compounds from the kinases and other enzymes totaled 18% (512). This is generally in agreement with the HEAL target distribution in the library and is reflective of the current status of pain and OUD targets being researched and developed. It also highlights the fact that most of the genome remains ‘undrugged’.

**Figure 1.**
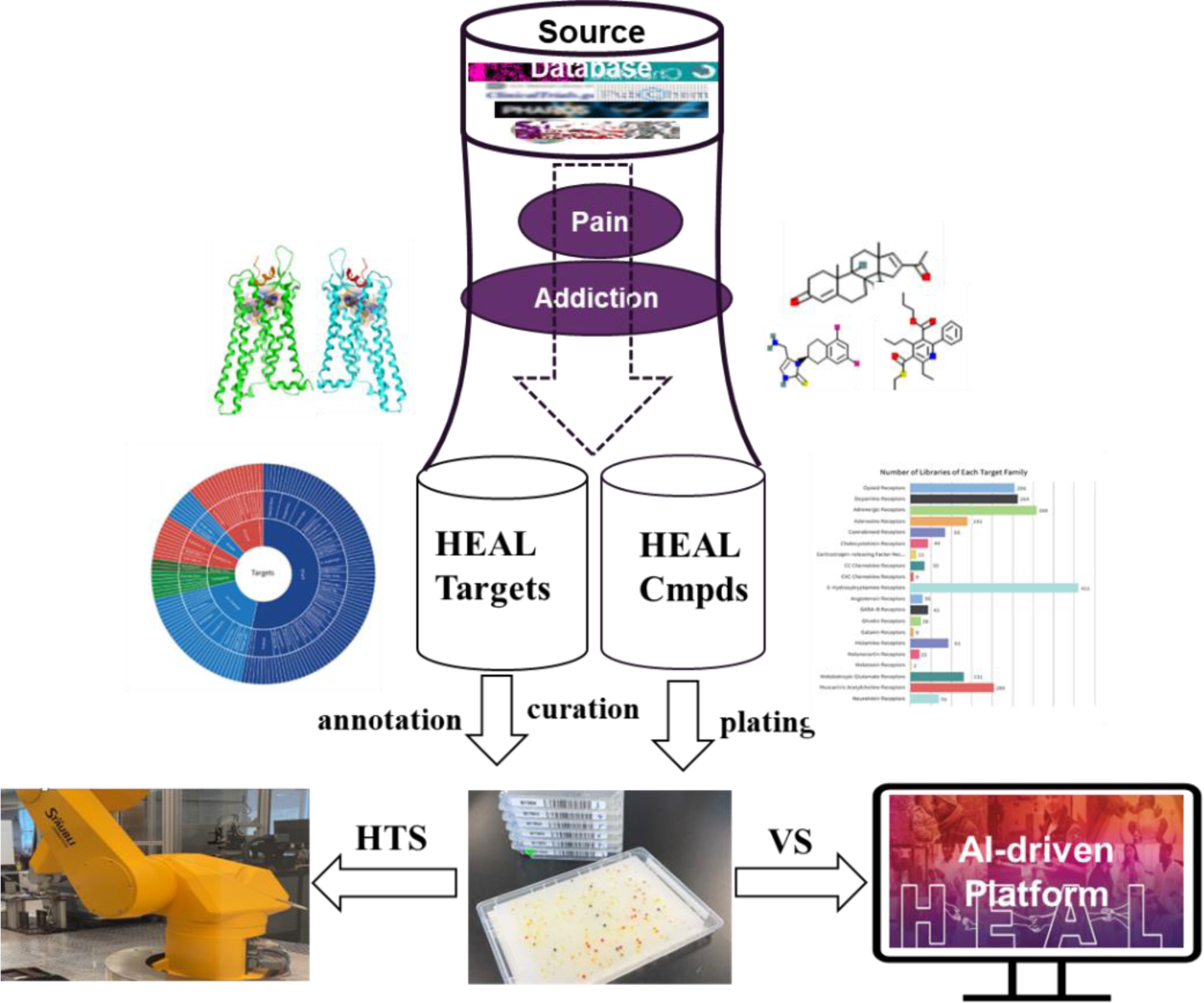
Collection of HEAL targets and compounds for quantitative high throughput screening and AI-driven drug discovery.

**Figure 2.**
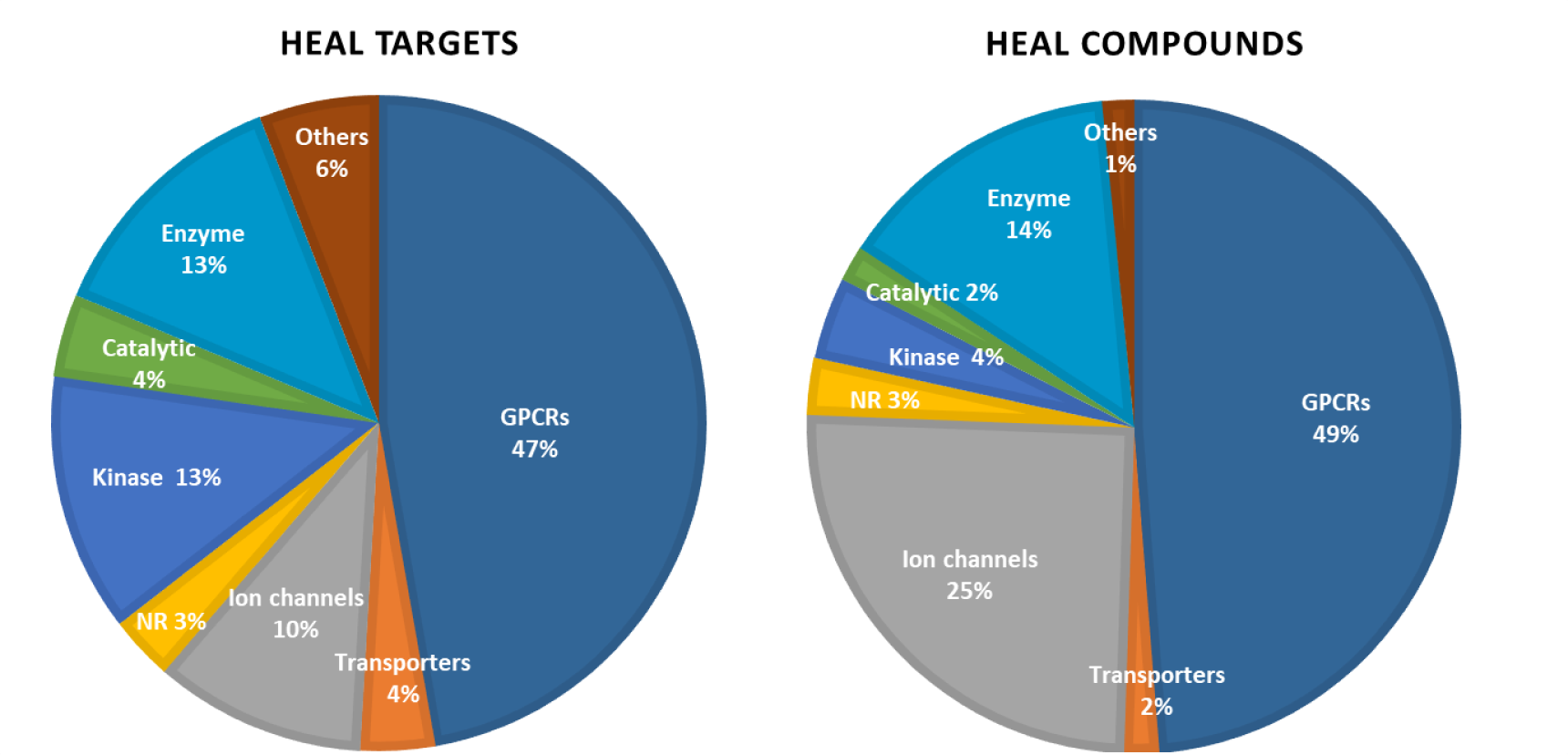
Components of HEAL targets and compounds.

We analyzed the physicochemical properties and structural features of the HEAL Library compounds. **Figure 3A** shows the overall distributions of molecular weight (MW), H-bond acceptor (HBA) and H-bond donor (HBD) counts, and topological polar surface area (TPSA) among these subsets of the HEAL compounds (clinical drugs, pain and additive drugs, CNS compounds, probes, and active compounds). Interestingly, the calculated CNS Multiparameter Optimization (MPO_ scores and Blood-Brain Barrier (BBB) permeability (**Figure 3B and 3C**) of HEAL compounds are generally “CNS-druglike” and comparable to the subset of pain-related drug compounds. Since the CNS property is a vital consideration for most pain and OUD drug development, the “CNS-likeness” of the HEAL Library also makes it a valuable dataset for a broad HTS screening as well as modeling against CNS-related diseases.

**Figure 3.**
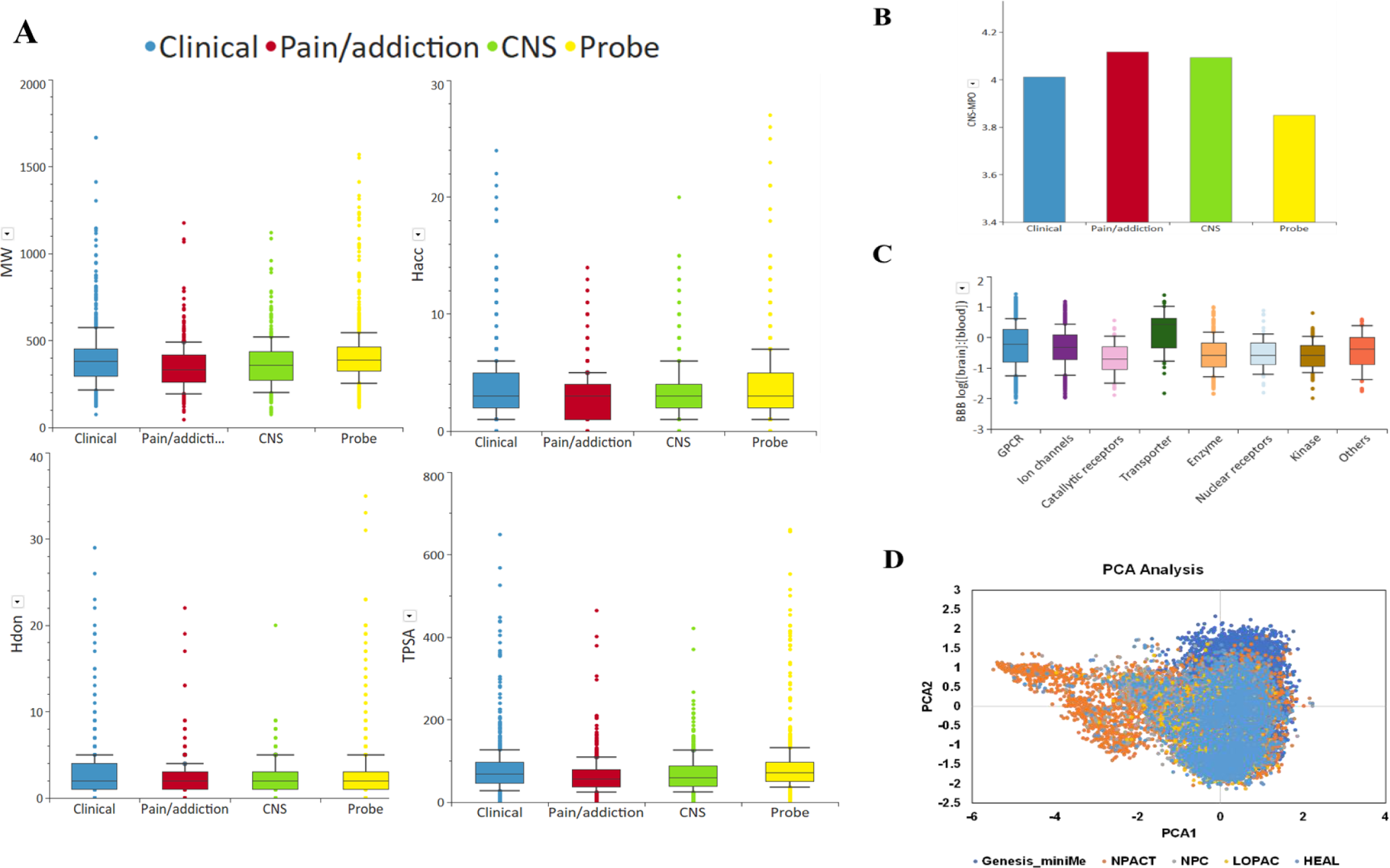
Physicochemical, CNS and BBB properties of HEAL library compounds.

We further compared the structural diversity and chemical space of the HEAL Library with other NCATS annotated libraries (**Figure 3D**). While the HEAL compounds mostly resembled the NPC approved drug collection, as aligned with Lipinki’s “rule of 5” ^28^, they also demonstrated an expansion into and overlap with the chemical diversity screening library (Genesis MiniMe collection), as well as an annotated library with a broad collection of pharmacologically active compounds and natural products (NPACT). The diversity of structural chemotypes and chemical space of HEAL compounds, therefore, provided a valuable reservoir as starting points for drug repurposing and rational drug design for novel pain and OUD targets.

## Compound plating for quantitative high throughput screening (qHTS)

Of 2,980 compounds collected, physical samples of 2,800 compounds were acquired. Except for a few controlled substances, all compounds were purchased from major chemical suppliers. The same compound acquisition process was used for creating and updating the NPC library ^12^. Once compounds were procured, quality control (QC) was performed by liquid chromatography-mass spectrometry (LC-MS) at NCATS. Compounds exhibiting less than 95% purity were purified further to achieve this minimum value of purity. Stock solutions were prepared at 10mL in DMSO after QC. All compounds were then plated in qHTS 1536-well plate format, both in acoustic source plates and pin tool plates for qHTS with 1:5 dilution and 7 concentrations, thus enabling a rapid and large-scale screening against a wide range of assays on the NCATS qHTS robot system.

## Profiling of HEAL Compounds for cell viability

We profiled the HEAL Library using a cell killing assay against the HeLa and HEK 293 cell lines to evaluate the cytotoxicity of the library’s compounds. HeLa is one of the commonly used human immortalized cell line, and the HEK 293 is a neuronal lineage line. Both are extensively employed in scientific research for pain and addiction studies. The HEAL Library compounds were tested with a 7-point dose-response CTG assay in these two cell lines. As shown in **Figure 4**, 20% and 24% of the library compounds were cytotoxic to HeLa and HEK 293 cells, respectively. Of these actives, 126 compounds overlapped and shared similar cytotoxic activity in both cell lines. Notably, the percentage of cytotoxic compounds within the HEAL Library for these two cell lines was generally lower than other annotated libraries we previously evaluated ^29^. Further analysis showed that the majority of the cytotoxic compounds were associated with kinase targets (this family is primarily comprised of probes and tool compounds). In contrast, cytotoxic compounds were less prevalent in the GPCR and other families. This result is anticipated and consistent with the promiscuity score of these drug compounds associated with these targets (**Figure 4D**).

**Figure 4.**
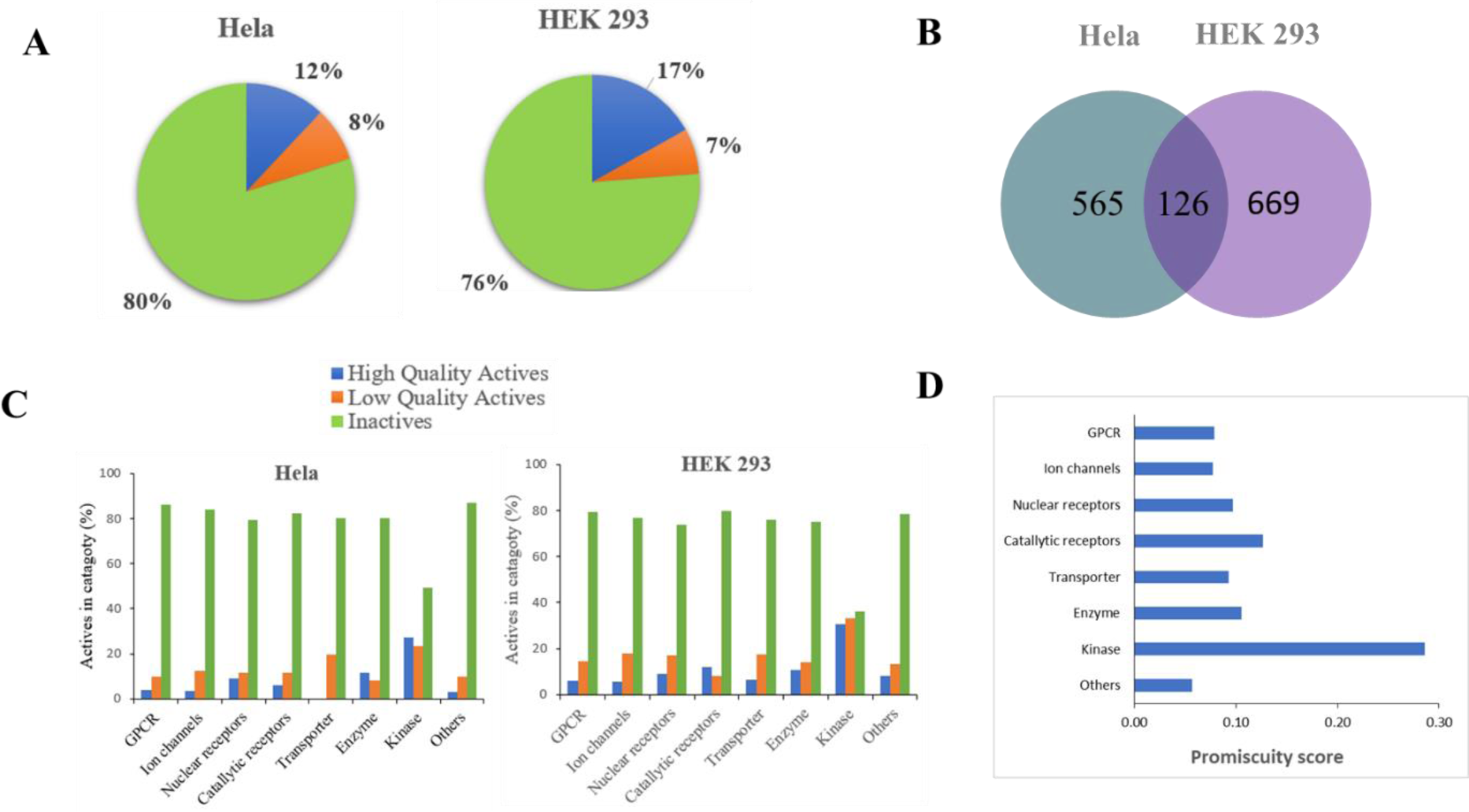
Profiling of HEAL library compounds A) cytotoxic compounds in Hella and HEK293 cells, B) overlap of cytotoxic compounds in HEK 293 and HeLa cells, C) actives in each target class, D) promiscuity scores.

## Application of HEAL library in qHTS/VS pilot screening

The HEAL Library has been utilized to screen against several HEAL targets as part of active collaborations between NCATS scientists and extramural scientists, including GAL1R, GPR151, PIEZO2, relaxin-3 receptor (RXFP3), the pituitary adenylate cyclase-activating polypeptide type I receptor (PAC1R), atrial natriuretic peptide receptor 1 (NPR1), adenyl cyclase 1 (AC1), and glutamate carboxypeptidase II (GCPII) (**Table 1**). Except for GCPII, a valid target for non-opioid drugs to treat pain (many potent but non-druggable small molecule inhibitors have been reported ^30^), very few modulators are known for these HEAL targets. Therefore, the HEAL Library provided a resource for pilot screening against these targets to identify potential active hits or tool compounds for further assay development and large-scale HTS.

All screens were conducted with various biochemical or cell-based assays, including counter assays developed for each target, all of which were miniaturized for qHTS at NCATS. Hits were determined based on the dose-response curve class as well as efficacy ^31^. The performance of hit rates in the primary screen across these 8 targets was shown in **Table 3**, which ranged from 0.15% - 6%. The hit rate associated with the two cyclase targets (NPR1 at 4.7% and AC1 at 6.0%) and PAC1R (5.7%) were relatively higher. Analysis showed that many of these “primary-assay-hits” may be non-selective or artifacts, suggesting that a secondary assay on their sister targets, for example, NPR2 and AC8, are needed to triage the hit list. The GCPII hits were mostly prior art compounds collected in the HEAL Library, indicating the robustness of the MS-LC assay used in the screen ^32^. The hits from the GAL1R and GPR151 screens showed some activity on both targets and were more appealing. GPR151 is also known as GALR4, sharing homology with the galanin receptor family ^33^. The small molecule agonists of GAL1R and GPR151 identified from the HEAL Library may provide novel candidates for further study.

**Table 3.** HEAL library screening against 8 targets using qHTS and VS. The VS enrichment factor is calculated by EF (%) = Act_sel_/ Act_tot_ x N_tot_/N_sel_. where Act_sel_ = the number of hits identified from VS; N_sel_ = the number of top-ranked compounds selected from VS (150 in this study); Act_tot_ = the total number of hits identified from HTS; N_tot_ = the total number of compounds screened (2800 in this study).

| Target | HTS |  | VS |  |
| --- | --- | --- | --- | --- |
|  | hits | hit rate (%) | hits | EF (%) |
| GAL1R | 4 | 0.14 | 1 | 4.67 |
| GPR151 | 18 | 0.64 | 5 | 5.19 |
| RXFP3 | 77 | 2.73 | 21 | 5.09 |
| PAC1R | 159 | 5.65 | 25 | 2.94 |
| PZ2 | 19 | 0.67 | 1 | 1.00 |
| GCPII | 26 | 0.92 | 11 | 7.90 |
| NPR1 | 132 | 4.69 | 17 | 2.40 |
| AC1 | 169 | 6.00 | 15 | 1.66 |

To further explore the qHTS hits from the HEAL library, we performed structure-based docking and virtual screening using an AI-driven platform integrated with structural modeling of the HEAL targets and compounds (**Supplementary Figure S1**). **Figure 5** shows the heatmap of VS hits in the HEAL Library in comparison with the hits from the experimental qHTS. The VS hit enrichment, which is defined as the qHTS hits pulled out from the top-ranked VS hitlist, varied from 1.0 (PZ2) to 7.94 (GCPII) (**Table 3**). This was in general agreement with the docking-based VS performance ^34^. The notably higher enrichment with GCPII can be attributed to its well-defined active site of crystal structures bound with various inhibitors, whereas for other targets (GPR151, AC1, NPR1), homology models were used in the docking. The ion channel PZ2 is a challenging target for assay development and screening, as well as structure-based VS. The 3D structures of PZ2 and PZ1 were solved recently, but the small molecule binding site remains elusive ^35, 36^. The predicted binding models of these potential hits with PZ2 may provide insight to probe the mechanism of action of ligand binding and activation.

**Figure 5.**
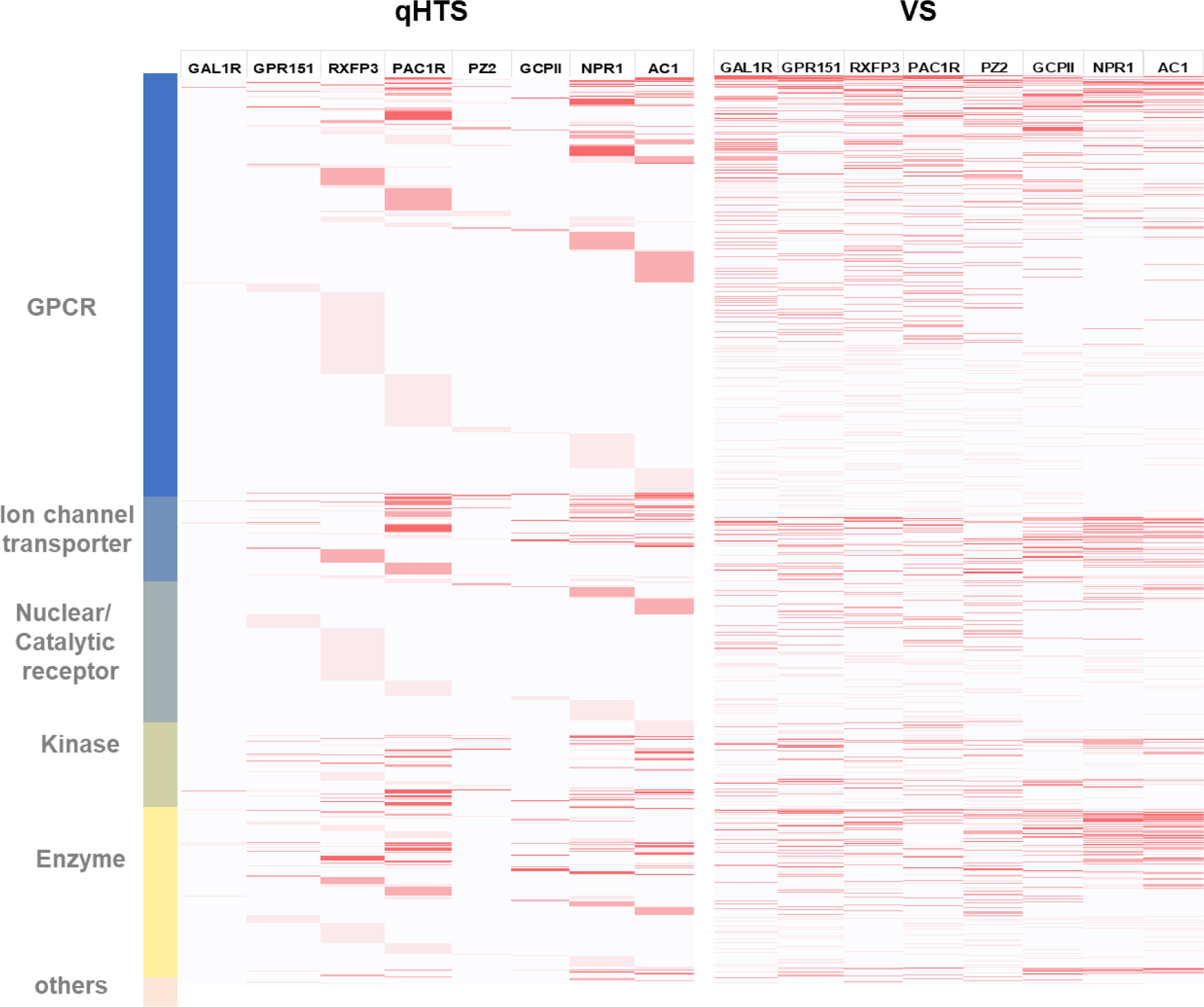
Heatmap of HEAL compound activity in qHTS and docking scores in VS on 8 HEAL targets.

Comparison of the qHTS and VS hits in the HEAL library pilot screening revealed some interesting findings. As a proof of concept, the results of both qHTS and VS showed multiple hits for each target that exceeded their pre-established category. The robust and potent qHTS-hits of the two GPCR targets RXFP3 and PAC1R were also found in the kinase and enzyme classes, while many potent hits of the two cyclase NPR1 and AC1 were found in the GPCR class (**Figure 5**). This indicated possible off-target effects due to the binding similarity at the active site of these target classes, and suggested an approach to triage the cross-targeted hits with a secondary assay such as GPCR or kinase profiling. On the other hand, the VS hits showed target enrichment in general. For example, the top VS-hits with these four GPCR targets were typically found in the GPCR class of the HEAL Library, and the top VS-hits for NPR1 and AC1 were mostly identified in the enzyme class. These results reflected the structural similarity of the binding pocket within the target family, which could be used for further qHTS-hit triage and lead prioritization.

## Outlook and Conclusions

While the HEAL Library plates are used routinely for HEAL project screening at NCATS, a pre-spotted copy of the HEAL Library in 384-well plate format (single concentration, 10mM) can be made available to the research community specifically for meritorious HEAL-related assays through NCATS Division of Preclinical Innovation (DPI) collaborations (https://ncats.nih.gov/research/research-activities/heal/expertise/library). Investigators interested in accessing the HEAL Library can submit a two-page proposal to NCATS DPI for evaluation. If selected, an overview of the applicant’s primary screening results will be shared and discussed with NCATS to determine whether spotted, cherry-picked plates of hits will be provided for counter screen assays. Plated copies of the HEAL Target and Compound Library as well as library cherry picks will only be provided as resources are available. All HEAL Library screening data are expected to be uploaded to PubChem within two years.

AI-driven machine learning and virtual screening are playing an ever-increasingly important role in for the field of rational drug discovery and development ^37^. With the accumulation of qHTS and VS data on the HEAL Library, we implemented and evaluated several noteworthy modules within a HEAL-focused AI-platform (**Figure 6**). For example, the 3D structures of all HEAL targets (Structure-DB) are collected from Protein Data Bank (PDB) or modeled by AlphaFold ^38^. The binding pockets of representative structures for each target and family are clustered and extracted (Pocket-DB). Systematic docking of all HEAL Library compounds to these binding pockets is performed, and the predicted binding models of each target-drug complex are deposited (Binding-DB). Additionally, ligand-based machine learning models for each target and family are generated based on their qHTS bioassay data (ML-DB).

**Figure 6.**
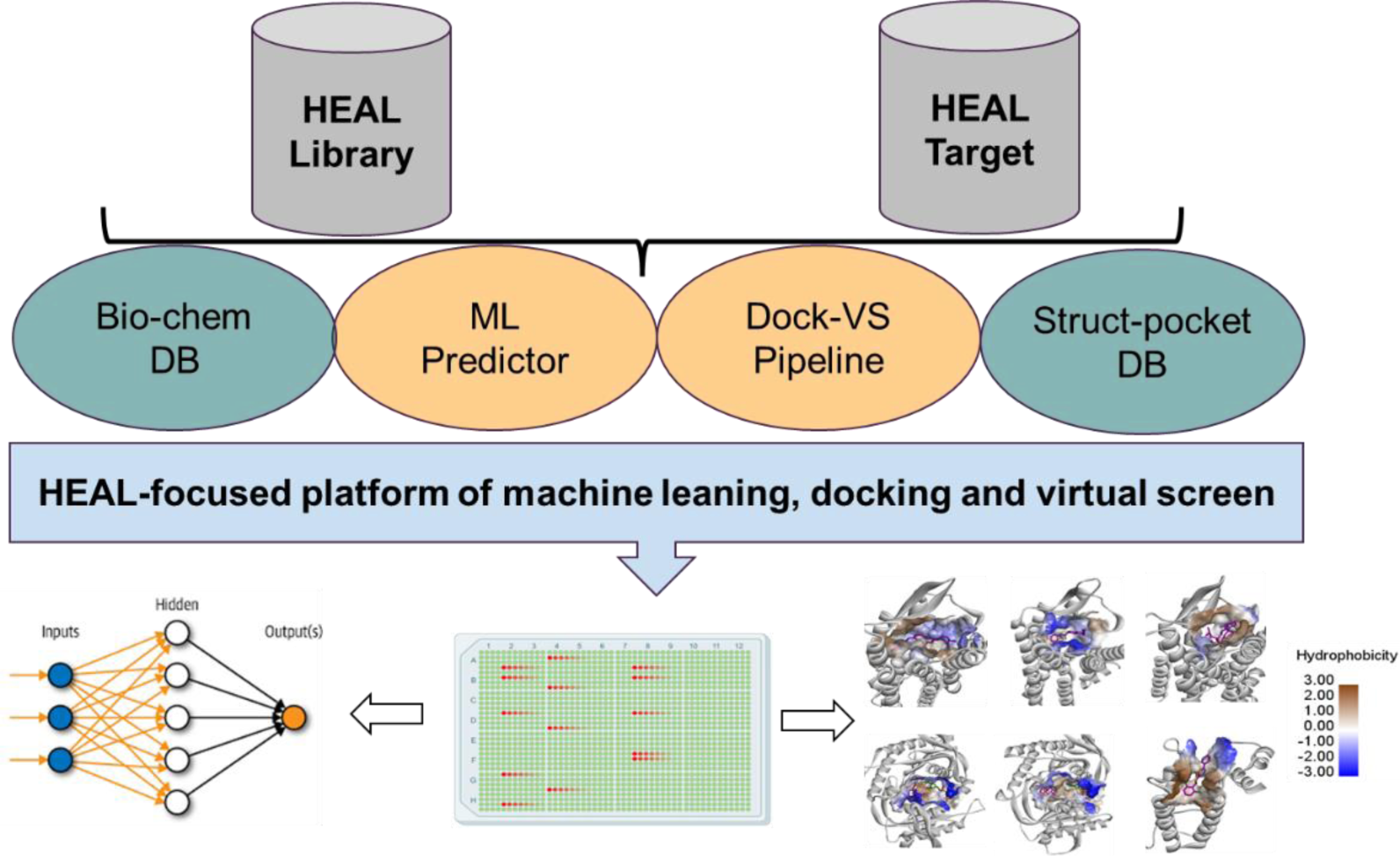
An AI-driven platform of HEAL compounds and targets for structure-based drug discovery.

The integration of these HEAL-focused modules, along with a qHTS open data portal ^39^, provides access to the abundant assay screening data generated from HEAL Library, and it facilitates efficient virtual screening of ultra-large chemical databases, such as Enamine’s REAL library containing billions of compounds.

In summary, the creation of the HEAL Library combined with an AI-driven platform provided a valuable resource to establish assay screening and collaborations, strategically advancing drug development programs for pain and OUD. A notable feature of the HEAL Library is the assay-ready pre-spotted plates which can be made available to extramural sites for HEAL-related assays through NCATS. Currently, the HEAL library has been screened against various HEAL targets and other relevant neurodegenerative projects. Similar to other NCATS annotated libraries like NPC and NPACT, the HEAL Library serves as a versatile tool for assay and target validation, exploration of mechanisms of action (MOA), as well as pilot screening of novel candidates. This not only enhances our understanding of the intricate processes involved but also paves the way for further development of probes and drugs aimed at treating pain and OUD.

## Supporting Information

List of annotated HEAL library compounds and target; Details of VS on 8 HEAL targets in the pilot screening.

## Supporting information

HEAL targets and compounds

## Acknowledgment

This research was supported in part by the Intramural research program of the National Center for Advancing Translational Sciences (National Institutes of Health, USA). Support was also provided by the Trans-NIH HEAL Initiative. The NIH HEAL Initiative Data policy is available here: https://heal.nih.gov/about/public-access-data.

## Notes

### Competing Interest Statement

The authors have declared no competing interest.

### Summary of Updates

Title revised; Figures revised; Tables revised.

