## Supplementary material for "A Comprehensive Collection of Pain and Opioid Use Disorder Compounds for High-Throughput Screening and Artificial Intelligence-Driven Drug Discovery": HEAL targets and compounds

**Table of Contents**


- **Figure S1:** Structural models of 8 HEAL targets in the pilot screening (Pg. S-3)
- **Table S1:** List of annotated HEAL library compounds and targets (Pg. S-4)

**Figure S1**. Structural models of 8 HEAL targets in the pilot screening. Protein surface at the active site is shown in hydrophobicity and small molecules in the binding pocket are shown in sticks. The structures of three available proteins were retrieved from the Protein Data Bank: GAL1R (7WQ3), PAC1R (6M1I), GCPII (4MCQ). The homology model of human PZ2 was generated from its homolog of mouse PZ2 structure (6KG7). The other four protein structural models were obtained from AlphaFold2 Protein Structure Database (https://alphafold.ebi.ac.uk/): GPR151 (UniProt ID: Q8TDV0), RXFP3 (Q9NSD7), AC1 (Q08828), NPR1 (P16066).

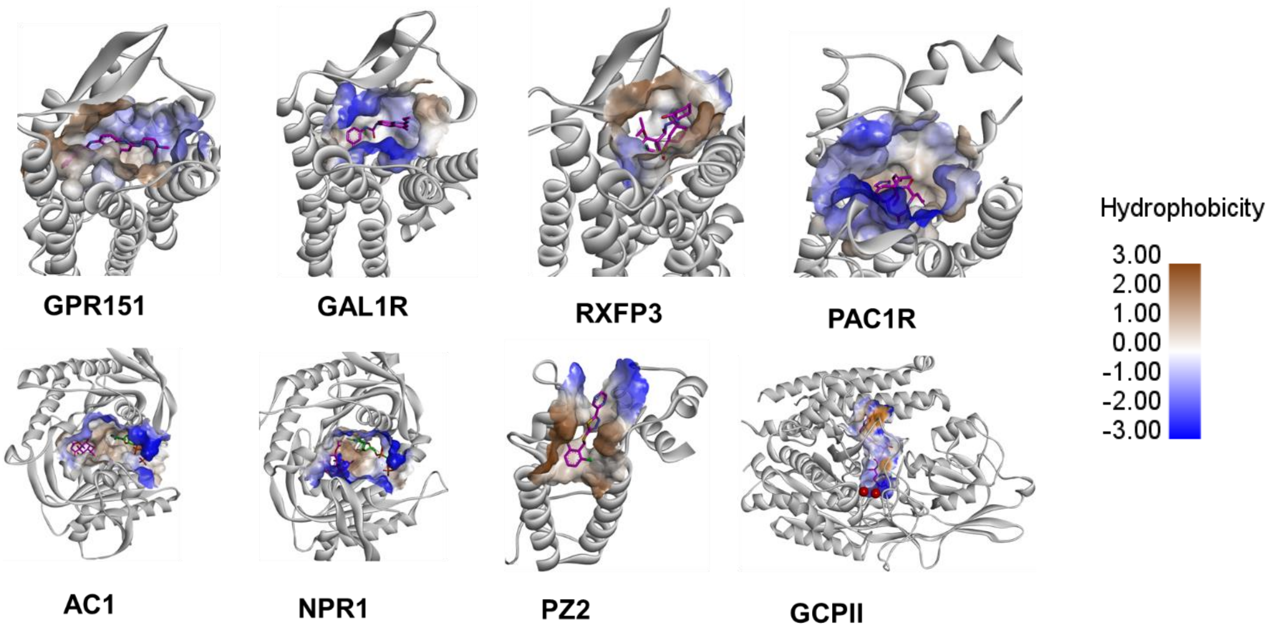

Table S1. List of annotated HEAL library compounds and targets.

| | | | | | | |
|---|---|---|---|---|---|---|
| NCGC00480731-01 | DREADD agonist 21 | mAChR | Muscarinic Acetylcholine Receptors | Agonist | Probe | GPCR |
| NCGC00379124-01 | SR 16584 | mAChR | Muscarinic Acetylcholine Receptors | Antagonist | Probe | GPCR |
| NCGC00167321-02 | Men 10376 | NK1R | Tachykinin receptor | Antagonist | Probe | GPCR |
| NCGC00167124-02 | Substance P (1-7) | NK1R | Tachykinin receptor | Agonist | Probe | GPCR |
| NCGC00685440-01 | Neurokinin A(4-10) | NK1R | Tachykinin receptor | Agonist | Probe | GPCR |
| NCGC00686680-01 | Senktide | NK1R | Tachykinin receptor | Agonist | Probe | GPCR |
| NCGC00378898-02 | NKP-608 | NK1R | Tachykinin receptor | Antagonist | Probe | GPCR |
| NCGC00159560-02 | FK-888 | NK1R | Tachykinin receptor | Antagonist | Probe | GPCR |
| NCGC00655648-01 | TAK-637 | NK1R | Tachykinin receptor | Antagonist | Probe | GPCR |
| NCGC00390587-04 | G-007-LK | NK1R | Tachykinin receptor | Antagonist | Probe | GPCR |
| NCGC00379214-07 | WIKI4 | NK1R | Tachykinin receptor | Antagonist | Probe | GPCR |
| NCGC00390606-02 | TNKS-656 | NK1R | Tachykinin receptor | Antagonist | Probe | GPCR |
| NCGC00387054-01 | L-732138 | NK1R | Tachykinin receptor | Antagonist | Probe | GPCR |
| NCGC00381706-01 | GR-159897 | NK1R | Tachykinin receptor | Antagonist | Probe | GPCR |
| NCGC00015956-07 | SB-222200 | NK1R | Tachykinin receptor | Antagonist | Probe | GPCR |
| NCGC00185684-04 | NCGC-84 | NPSR | Neuropeptide S | Antagonist | Probe | GPCR |
| NCGC00189027-01 | QA1 | NPSR | Neuropeptide S | Antagonist | Probe | GPCR |
| NCGC00184846-02 | SHA-68 | NPSR | Neuropeptide S | Antagonist | Probe | GPCR |
| NCGC00378699-01 | BIBP-3226 | NPY1R | Neuropeptide Y receptor Y1 | Antagonist | Probe | GPCR |
| NCGC00371076-01 | BIBO3304 | NPY1R | Neuropeptide Y receptor Y1 | Antagonist | Probe | GPCR |
| NCGC00485144-03 | BMS-193885 | NPYR | Neuropeptide Y receptor | Antagonist | Probe | GPCR |
| NCGC00390574-03 | JNJ-31020028 | NPYR | Neuropeptide Y receptor | Antagonist | Probe | GPCR |
| NCGC00371082-02 | CGP-71683 | NPYR | Neuropeptide Y receptor | Antagonist | Probe | GPCR |
| NCGC00371015-01 | JNJ-5207787 | NPYR | Neuropeptide Y receptor | Antagonist | Probe | GPCR |
| NCGC00379167-01 | CYM-9484 | NPYR | Neuropeptide Y receptor | Antagonist | Probe | GPCR |
| NCGC00485847-02 | RF9 | NPYR | Neuropeptide Y receptor | Antagonist | Probe | GPCR |
| NCGC00370968-02 | SF-11 | NPYR | Neuropeptide Y receptor | Antagonist | Probe | GPCR |
| NCGC00387447-02 | GW-438014A | NPYR | Neuropeptide Y receptor | Antagonist | Probe | GPCR |
| NCGC00370875-02 | LU-AA33810 | NPYR | Neuropeptide Y receptor | Antagonist | Probe | GPCR |
| NCGC00370914-01 | NPY 5RA972 | NPYR | Neuropeptide Y receptor | Antagonist | Probe | GPCR |
| NCGC00485887-01 | AF38469 | NTR | Neurotensin Receptors | Antagonist | Probe | GPCR |
| NCGC00378708-01 | ML314 | NTR1 | Neurotensin receptor type 1 | Agonist | Probe | GPCR |
| NCGC00485321-01 | Naltrindole isothiocyanate | DOR | delta receptor | Antagonist | Probe | GPCR |
| NCGC00485085-01 | Butorphan | DOR | delta receptor | Agonist | Probe | GPCR |
| NCGC00379218-03 | ML190 | KOR | kappa receptor | Antagonist | Probe | GPCR |
| NCGC00163188-01 | DSLET | MOR | mu receptor | Agonist | Probe | GPCR |
| NCGC00487379-01 | BMS-986124 | MOR | mu receptor | NAM | Probe | GPCR |
| NCGC00408804-01 | LY-255582 | MOR | mu receptor | Antagonist | Probe | GPCR |
| NCGC00485892-01 | 6-Alpha Naloxol | MOR | mu receptor | Antagonist | Probe | GPCR |
| NCGC00685453-01 | LY-2940094 | NOPR | Nociceptin receptor | Antagonist | Probe | GPCR |
| NCGC00387110-03 | C24 | NOPR | Nociceptin receptor | Antagonist | Probe | GPCR |
| NCGC00346896-01 | MCOPPB | NOPR | Nociceptin receptor | Agonist | Probe | GPCR |
| NCGC00655635-01 | Ro 65-6570 | NOPR | Nociceptin receptor | Agonist | Probe | GPCR |
| NCGC00389733-01 | Adrenorphin | OR | Opioid receptors | Agonist | Probe | GPCR |

| | | | | | | |
|---|---|---|---|---|---|---|
| NCGC00685434-01 | SR-17018 | OR | Opioid receptors | Agonist | Probe | GPCR |
| NCGC00386865-02 | ML-335 | OR | Opioid receptors | Agonist | Probe | GPCR |
| NCGC00601700-01 | LY-2795050 | OR | Opioid receptors | Antagonist | Probe | GPCR |
| NCGC00189140-06 | UNC0642 | Sigma1R | sigma 1-type opioid receptor | Antagonist | Probe | GPCR |
| NCGC00024766-03 | 4-IBP | Sigma1R | sigma 1-type opioid receptor | Agonist | Probe | GPCR |
| NCGC00165875-03 | PB-28 | Sigma1R | sigma 1-type opioid receptor | Agonist | Probe | GPCR |
| NCGC00015940-05 | SKF-83959 | Sigma1R | sigma 1-type opioid receptor | Partial agonist | Probe | GPCR |
| NCGC00387082-01 | NE-100 | Sigma1R | sigma 1-type opioid receptor | Antagonist | Probe | GPCR |
| NCGC00371106-01 | PPCC | Sigma1R | sigma 1-type opioid receptor | Agonist | Probe | GPCR |
| NCGC00024844-04 | BD-1063 | Sigma1R | sigma 1-type opioid receptor | Antagonist | Probe | GPCR |
| NCGC00378668-01 | PD 144418 oxalate | Sigma1R | sigma 1-type opioid receptor | Antagonist | Probe | GPCR |
| NCGC00024819-02 | Ditolylguanidine | Sigma1R | sigma 1-type opioid receptor | Agonist | Probe | GPCR |
| NCGC00482938-01 | Schisantherin A | OX | Hypocretin receptor (HCRTR) | Antagonist | Probe | GPCR |
| NCGC00379105-02 | JNJ-10397049 | OX | Hypocretin receptor (HCRTR) | Antagonist | Probe | GPCR |
| NCGC00685423-01 | SB-649868 | OX | Hypocretin receptor (HCRTR) | Antagonist | Probe | GPCR |
| NCGC00370975-03 | TCS-1102 | OX | Hypocretin receptor (HCRTR) | Antagonist | Probe | GPCR |
| NCGC00510490-03 | MK-1064 | OX | Hypocretin receptor (HCRTR) | Antagonist | Probe | GPCR |
| NCGC00386678-01 | EMPA | OX | Hypocretin receptor (HCRTR) | Antagonist | Probe | GPCR |
| NCGC00685393-01 | Nemorexant | OX | Hypocretin receptor (HCRTR) | Antagonist | Probe | GPCR |
| NCGC00379073-04 | SB-674042 | OX | Hypocretin receptor (HCRTR) | Antagonist | Probe | GPCR |
| NCGC00387851-03 | MK-3697 | OX | Hypocretin receptor (HCRTR) | Antagonist | Probe | GPCR |
| NCGC00402232-03 | GSK-1059865 | OX | Hypocretin receptor (HCRTR) | Antagonist | Probe | GPCR |
| NCGC00370876-02 | TCS-OX2-29 | OX | Hypocretin receptor (HCRTR) | Antagonist | Probe | GPCR |
| NCGC00387479-01 | ACT-462206 | OX | Hypocretin receptor (HCRTR) | Antagonist | Probe | GPCR |
| NCGC00508857-03 | VC-5220 | OX1 | Orexin receptor type 1 | Agonist | Probe | GPCR |
| NCGC00370769-01 | ACT-335827 | OX1 | Orexin receptor type 1 | Antagonist | Probe | GPCR |
| NCGC00015602-05 | L-368899 | OT | Oxytocin receptor | Antagonist | Probe | GPCR |
| NCGC00159562-02 | L-371257 | OT | Oxytocin receptor | Antagonist | Probe | GPCR |
| NCGC00481601-02 | Cligosiban | OT | Oxytocin receptor | Antagonist | Probe | GPCR |
| NCGC00389333-01 | Alprostadil | EP1 | prostaglandin E1 receptor | Antagonist | Probe | GPCR |
| NCGC00476132-02 | MRE-269 | IP | Prostaglandin I2 (IP) receptor | inhibitor | Probe | GPCR |
| NCGC00250135-09 | ML290 | RXFP1 | Relaxin family peptide receptor 1 | Agonist | Probe | GPCR |
| NCGC00014994-10 | Tryptamine | TAAR1 | Trace amine receptor | Agonist | Probe | GPCR |
| NCGC00182033-01 | Felypressin | V1AR | Vasopressin V1A rec; Antidiuretic hormone receptor 1A | Agonist | Probe | GPCR |
| NCGC00185754-02 | Terlipressin | V1BR | Vasopressin V1b rec; Antidiuretic hormone receptor 1b \| AVPR3 \| V3; | Agonist | Probe | GPCR |
| NCGC00181743-02 | Lypressin | V1BR | Vasopressin V1b rec; Antidiuretic hormone receptor 1b \| AVPR3 \| V3; | Agonist | Probe | GPCR |
| NCGC00370961-02 | OPC-21268 | V1BR | Vasopressin V1b rec; Antidiuretic hormone receptor 1b \| AVPR3 \| V3; | Antagonist | Probe | GPCR |
| NCGC00423683-01 | PA-9 | PAC1R | Pituitary adenylate cyclase-activating polypeptide type I receptor | Antagonist | Probe | GPCR |
| NCGC00482524-01 | Jujuboside A | GABAA | GABAA receptor | Agonist | Probe | Ion channel |
| NCGC00094227-07 | Picrotoxin | GABAA | GABAA receptor | Antagonist | Probe | Ion channel |
| NCGC00686691-01 | Cholesterol myristate | GABAA | GABAA receptor | Antagonist | Probe | Ion channel |

| | | | | | | |
|---|---|---|---|---|---|---|
| NCGC00686693-01 | Fluxametamide | GABAA | GABAA receptor | Antagonist | Probe | Ion channel |
| NCGC00386589-01 | CP-615003 | GABAA | GABAA receptor | Partial agonist | Probe | Ion channel |
| NCGC00024989-03 | CGP-54626 | GABAA | GABAA receptor | Antagonist | Probe | Ion channel |
| NCGC00379135-01 | PHP-501 | GABAA | GABAA receptor | Antagonist | Probe | Ion channel |
| NCGC00161395-02 | ZK-93423 | GABAA | GABAA receptor | Agonist | Probe | Ion channel |
| NCGC00025245-04 | Ginkgolide B | GABAA | GABAA receptor | Antagonist | Probe | Ion channel |
| NCGC00685337-01 | Zuranolone | GABAA | GABAA receptor | Agonist | Probe | Ion channel |
| NCGC00370896-02 | L-838417 | GABAA | GABAA receptor | Agonist | Probe | Ion channel |
| NCGC00402272-01 | Imidazenil | GABAA | GABAA receptor | PAM | Probe | Ion channel |
| NCGC00510096-02 | DAA-1106 | GABAA | GABAA receptor | Agonist | Probe | Ion channel |
| NCGC00685444-01 | SSD-114 | GABAA | GABAA receptor | PAM | Probe | Ion channel |
| NCGC00025075-03 | CGP-52432 | GABAA | GABAA receptor | Antagonist | Probe | Ion channel |
| NCGC00685351-01 | ONO-8590580 | GABAA | GABAA receptor | NAM | Probe | Ion channel |
| NCGC00015977-06 | SKF-89976A | GABAA | GABAA receptor | Antagonist | Probe | Ion channel |
| NCGC00015909-14 | Gabazine | GABAA | GABAA receptor | Antagonist | Probe | Ion channel |
| NCGC00487171-01 | alpha5IA | GABAA | GABAA receptor | Inverse agonist | Probe | Ion channel |
| NCGC00485016-01 | Hydroxytriazolam | GABAA | GABAA receptor | Agonist | Probe | Ion channel |
| NCGC00386556-02 | NS-11394 | GABAA | GABAA receptor | PAM | Probe | Ion channel |
| NCGC00378706-01 | U-90042 | GABAA | GABAA receptor | Agonist | Probe | Ion channel |
| NCGC00015392-06 | DMCM | GABAA | GABAA receptor | Inverse agonist | Probe | Ion channel |
| NCGC00655637-01 | alpha3IA | GABAA | GABAA receptor | Inverse agonist | Probe | Ion channel |
| NCGC00161396-02 | ZK-93426 | GABAA | GABAA receptor | Antagonist | Probe | Ion channel |
| NCGC00015621-02 | L-655708 | GABAA | GABAA receptor | Inverse agonist | Probe | Ion channel |
| NCGC00370965-01 | TCS-1205 | GABAA | GABAA receptor | Partial agonist | Probe | Ion channel |
| NCGC00510918-01 | Progabide | GABAA | GABAA receptor | Agonist | Probe | Ion channel |
| NCGC00163288-01 | Tetrahydrodeoxycorticosterone | GABAA | GABAA receptor | Agonist | Probe | Ion channel |
| NCGC00092292-02 | Ro 15-4513 | GABAA | GABAA receptor | Inverse agonist | Probe | Ion channel |
| NCGC00025337-02 | Ro 19-4603 | GABAA | GABAA receptor | Inverse agonist | Probe | Ion channel |
| NCGC00378712-02 | U-93631 | GABAA | GABAA receptor | Agonist | Probe | Ion channel |
| NCGC00370973-01 | TCS-1105 | GABAA | GABAA receptor | Antagonist | Probe | Ion channel |
| NCGC00408906-01 | CGS-9895 | GABAA | GABAA receptor | Antagonist | Probe | Ion channel |
| NCGC00015077-06 | 2-Hydroxysaclofen | GABAA | GABAA receptor | Antagonist | Probe | Ion channel |
| NCGC00599617-01 | CGS-8216 | GABAA | GABAA receptor | Antagonist | Probe | Ion channel |
| NCGC00389924-01 | 6,2'-Dihydroxyflavone | GABAA | GABAA receptor | Antagonist | Probe | Ion channel |
| NCGC00024514-04 | Saclofen | GABAA | GABAA receptor | Antagonist | Probe | Ion channel |
| NCGC00015781-05 | Phaclofen | GABAA | GABAA receptor | Antagonist | Probe | Ion channel |
| NCGC00095563-04 | TMCA | GABAA | GABAA receptor | Antagonist | Probe | Ion channel |
| NCGC00142612-02 | 6-Methylflavone | GABAA | GABAA receptor | Agonist | Probe | Ion channel |
| NCGC00015391-08 | FG 7142 | GABAA | GABAA receptor | Inverse agonist | Probe | Ion channel |
| NCGC00025074-03 | CGP-35348 | GABAA | GABAA receptor | Antagonist | Probe | Ion channel |
| NCGC00025076-03 | CGP-46381 | GABAA | GABAA receptor | Antagonist | Probe | Ion channel |
| NCGC00015456-06 | Isoguvacine | GABAA | GABAA receptor | Agonist | Probe | Ion channel |
| NCGC00414969-02 | N-acetyl GABA | GABAA | GABAA receptor | Agonist | Probe | Ion channel |

| | | | | | | |
|---|---|---|---|---|---|---|
| NCGC00015015-04 | 3-APMPA | GABAA | GABAA receptor | Agonist | Probe | Ion channel |
| NCGC00188243-04 | p-Hydroxybenzaldehyde | GABAA | GABAA receptor | Antagonist | Probe | Ion channel |
| NCGC00013478-05 | L-DABA | GABAA | GABAA receptor | Antagonist | Probe | Ion channel |
| NCGC00371073-01 | CP-465022 | AMPA | GluA1; GluA2; GluA3 and GluA4 subunits | Antagonist | Probe | Ion channel |
| NCGC00485405-01 | PF-04725379 | AMPA | GluA1; GluA2; GluA3 and GluA4 subunits | PAM | Probe | Ion channel |
| NCGC00015379-05 | PEPA | AMPA | GluA1; GluA2; GluA3 and GluA4 subunits | PAM | Probe | Ion channel |
| NCGC00370740-02 | LY-450108 | AMPA | GluA1; GluA2; GluA3 and GluA4 subunits | PAM | Probe | Ion channel |
| NCGC00387210-02 | PF-4778574 | AMPA | GluA1; GluA2; GluA3 and GluA4 subunits | PAM | Probe | Ion channel |
| NCGC00263113-02 | GYKI-53655 | AMPA | GluA1; GluA2; GluA3 and GluA4 subunits | NAM | Probe | Ion channel |
| NCGC00387074-02 | Naspm | AMPA | GluA1; GluA2; GluA3 and GluA4 subunits | Antagonist | Probe | Ion channel |
| NCGC00025079-02 | SDZ 220-581 | AMPA | GluA1; GluA2; GluA3 and GluA4 subunits | Antagonist | Probe | Ion channel |
| NCGC00485396-01 | PF-04701475 | AMPA | GluA1; GluA2; GluA3 and GluA4 subunits | PAM | Probe | Ion channel |
| NCGC00370739-02 | LY-404187 | AMPA | GluA1; GluA2; GluA3 and GluA4 subunits | PAM | Probe | Ion channel |
| NCGC00387146-01 | IEM-1925 | AMPA | GluA1; GluA2; GluA3 and GluA4 subunits | Antagonist | Probe | Ion channel |
| NCGC00015093-05 | ATPO | AMPA | GluA1; GluA2; GluA3 and GluA4 subunits | Antagonist | Probe | Ion channel |
| NCGC00024988-03 | CFM-2 | AMPA | GluA1; GluA2; GluA3 and GluA4 subunits | Antagonist | Probe | Ion channel |
| NCGC00386525-02 | YM-90K | AMPA | GluA1; GluA2; GluA3 and GluA4 subunits | Antagonist | Probe | Ion channel |
| NCGC00015463-06 | GYKI-52466 | AMPA | GluA1; GluA2; GluA3 and GluA4 subunits | Antagonist | Probe | Ion channel |
| NCGC00024528-03 | 5-fluorowillardiine | AMPA | GluA1; GluA2; GluA3 and GluA4 subunits | Agonist | Probe | Ion channel |
| NCGC00015350-05 | Sunifiram | AMPA | GluA1; GluA2; GluA3 and GluA4 subunits | Antagonist | Probe | Ion channel |
| NCGC00025050-02 | IDRA-21 | AMPA | GluA1; GluA2; GluA3 and GluA4 subunits | PAM | Probe | Ion channel |
| NCGC00522557-01 | Org-26576 | AMPA | GluA1; GluA2; GluA3 and GluA4 subunits | PAM | Probe | Ion channel |
| NCGC00094464-04 | Willardiine | AMPA | GluA1; GluA2; GluA3 and GluA4 subunits | Agonist | Probe | Ion channel |
| NCGC00024517-03 | AMPA | AMPA | GluA1; GluA2; GluA3 and GluA4 subunits | Agonist | Probe | Ion channel |
| NCGC00378721-02 | Philanthotoxin 74 | iGluR | Ionotropic Glutamate Receptors | Antagonist | Probe | Ion channel |
| NCGC00370786-02 | CIQ | iGluR | Ionotropic Glutamate Receptors | Agonist | Probe | Ion channel |
| NCGC00025226-04 | Ro 25-6981 | iGLuR | Ionotropic Glutamate Receptors | Antagonist | Probe | Ion channel |
| NCGC00685441-01 | Neu2000 | iGluR | Ionotropic Glutamate Receptors | Antagonist | Probe | Ion channel |
| NCGC00685353-01 | Tulrampator | iGluR | Ionotropic Glutamate Receptors | PAM | Probe | Ion channel |
| NCGC00024904-03 | SYM2206 | iGluR | Ionotropic Glutamate Receptors | Antagonist | Probe | Ion channel |
| NCGC00024864-03 | L-701324 | iGluR | Ionotropic Glutamate Receptors | Antagonist | Probe | Ion channel |
| NCGC00015200-07 | CX-546 | iGluR | Ionotropic Glutamate Receptors | Agonist | Probe | Ion channel |
| NCGC00024475-05 | gamma-DGG | iGluR | Ionotropic Glutamate Receptors | Antagonist | Probe | Ion channel |

| | | | | | | |
|---|---|---|---|---|---|---|
| NCGC00650294-01 | 6-Methoxy-2-naphthoic acid | iGluR | Ionotropic Glutamate Receptors | PAM | Probe | Ion channel |
| NCGC00475818-01 | Ibotenic acid | iGluR | Ionotropic Glutamate Receptors | Agonist | Probe | Ion channel |
| NCGC00378702-01 | ACET | kainate | GluK1; GluK2 or GluK3 | Antagonist | Probe | Ion channel |
| NCGC00092326-03 | NS-3763 | Kainate | GluK1; GluK2 or GluK3 | Antagonist | Probe | Ion channel |
| NCGC00371089-01 | GYKI-47261 | kainate | GluK1; GluK2 or GluK3 | Antagonist | Probe | Ion channel |
| NCGC00370889-01 | UBP-310 | kainate | GluK1; GluK2 or GluK3 | Antagonist | Probe | Ion channel |
| NCGC00025352-02 | UBP-302 | Kainate | GluK1; GluK2 or GluK3 | Antagonist | Probe | Ion channel |
| NCGC00024529-03 | 5-iodowillardiine | Kainate | GluK1; GluK2 or GluK3 | Agonist | Probe | Ion channel |
| NCGC00024520-02 | Domoic acid | Kainate | GluK1; GluK2 or GluK3 | Agonist | Probe | Ion channel |
| NCGC00025003-02 | ATPA | Kainate | GluK1; GluK2 or GluK3 | Agonist | Probe | Ion channel |
| NCGC00024860-03 | SYM 2081 | Kainate | GluK1; GluK2 or GluK3 | Antagonist | Probe | Ion channel |
| NCGC00346887-05 | TCN-201 | NMDA | GluN1; GluN2A; GluN2B; GluN2C; GluN2D; GluN3A and GluN3B subunits | Antagonist | Probe | Ion channel |
| NCGC00485422-01 | PEAQX | NMDA | GluN1; GluN2A; GluN2B; GluN2C; GluN2D; GluN3A and GluN3B subunits | Antagonist | Probe | Ion channel |
| NCGC00379207-03 | QNZ46 | NMDA | GluN1; GluN2A; GluN2B; GluN2C; GluN2D; GluN3A and GluN3B subunits | Antagonist | Probe | Ion channel |
| NCGC00386738-01 | ZD-9379 | NMDA | GluN1; GluN2A; GluN2B; GluN2C; GluN2D; GluN3A and GluN3B subunits | Antagonist | Probe | Ion channel |
| NCGC00024761-03 | L-689560 | NMDA | GluN1; GluN2A; GluN2B; GluN2C; GluN2D; GluN3A and GluN3B subunits | Antagonist | Probe | Ion channel |
| NCGC00015639-06 | MDL-105519 | NMDA | GluN1; GluN2A; GluN2B; GluN2C; GluN2D; GluN3A and GluN3B subunits | Antagonist | Probe | Ion channel |
| NCGC00379098-01 | TCN-237 | NMDA | GluN1; GluN2A; GluN2B; GluN2C; GluN2D; GluN3A and GluN3B subunits | Antagonist | Probe | Ion channel |
| NCGC00378727-01 | TCS-46b | NMDA | GluN1; GluN2A; GluN2B; GluN2C; GluN2D; GluN3A and GluN3B subunits | Antagonist | Probe | Ion channel |
| NCGC00163269-04 | MDL-29951 | NMDA | GluN1; GluN2A; GluN2B; GluN2C; GluN2D; GluN3A and GluN3B subunits | Antagonist | Probe | Ion channel |
| NCGC00015313-06 | DCKA | NMDA | GluN1; GluN2A; GluN2B; GluN2C; GluN2D; GluN3A and GluN3B subunits | Antagonist | Probe | Ion channel |
| NCGC00015179-06 | CPP | NMDA | GluN1; GluN2A; GluN2B; GluN2C; GluN2D; GluN3A and GluN3B subunits | Antagonist | Probe | Ion channel |
| NCGC00485118-01 | 1-N-Methylamino-3,5-dimethyladamantane | NMDA | GluN1; GluN2A; GluN2B; GluN2C; GluN2D; GluN3A and GluN3B subunits | Antagonist | Probe | Ion channel |
| NCGC00025070-03 | Selfotel | NMDA | GluN1; GluN2A; GluN2B; GluN2C; GluN2D; GluN3A and GluN3B subunits | Antagonist | Probe | Ion channel |
| NCGC00486067-01 | LY-233053 | NMDA | GluN1; GluN2A; GluN2B; GluN2C; GluN2D; GluN3A and GluN3B subunits | Antagonist | Probe | Ion channel |
| NCGC00025174-02 | CGP-37849 | NMDA | GluN1; GluN2A; GluN2B; GluN2C; GluN2D; GluN3A and GluN3B subunits | Antagonist | Probe | Ion channel |
| NCGC00024492-03 | Homoquinolinic acid | NMDA | GluN1; GluN2A; GluN2B; GluN2C; GluN2D; GluN3A and GluN3B subunits | Partial agonist | Probe | Ion channel |

| | | | | | | |
|---|---|---|---|---|---|---|
| NCGC00024531-02 | Tetrazolylglycine | NMDA | GluN1; GluN2A; GluN2B; GluN2C; GluN2D; GluN3A and GluN3B subunits | Full agonist | Probe | Ion channel |
| NCGC00015111-02 | HA-966 | NMDA | GluN1; GluN2A; GluN2B; GluN2C; GluN2D; GluN3A and GluN3B subunits | Antagonist | Probe | Ion channel |
| NCGC00685390-01 | Methyllycaconitine | nAChR | Cholinergic receptor | Antagonist | Probe | Ion channel |
| NCGC00387214-03 | EG00229 | nAChR | Cholinergic receptor | Antagonist | Probe | Ion channel |
| NCGC00685363-01 | B-973B | nAChR | Cholinergic receptor | PAM | Probe | Ion channel |
| NCGC00387153-01 | 4BP-TQS | nAChR | Cholinergic receptor | Agonist | Probe | Ion channel |
| NCGC00371047-03 | A 867744 | nAChR | Cholinergic receptor | Agonist | Probe | Ion channel |
| NCGC00370848-01 | Desformylflustrabromine | nAChR | Cholinergic receptor | Agonist | Probe | Ion channel |
| NCGC00185993-02 | DHβE | nAChR | Cholinergic receptor | Antagonist | Probe | Ion channel |
| NCGC00487104-01 | GST-21 | nAChR | Cholinergic receptor | Agonist | Probe | Ion channel |
| NCGC00379128-01 | SEN 12333 | nAChR | Cholinergic receptor | Agonist | Probe | Ion channel |
| NCGC00015758-05 | Oxotremorine M | nAChR | Cholinergic receptor | Agonist | Probe | Ion channel |
| NCGC00371020-01 | A 844606 | nAChR | Cholinergic receptor | Agonist | Probe | Ion channel |
| NCGC00092364-11 | PNU-282987 | nAChR | Cholinergic receptor | Agonist | Probe | Ion channel |
| NCGC00378710-01 | Sazetidine A dihydrochloride | nAChR | Cholinergic receptor | Agonist | Probe | Ion channel |
| NCGC00370937-01 | A 582941 | nAChR | Cholinergic receptor | Agonist | Probe | Ion channel |
| NCGC00015370-03 | DBO-83 | nAChR | Cholinergic receptor | Agonist | Probe | Ion channel |
| NCGC00485128-01 | DBO-83 | nAChR | Cholinergic receptor | Agonist | Probe | Ion channel |
| NCGC00024730-02 | Epibatidine | nAChR | Cholinergic receptor | Agonist | Probe | Ion channel |
| NCGC00378539-01 | RJR 2429 | nAChR | Cholinergic receptor | Agonist | Probe | Ion channel |
| NCGC00387223-01 | A 85380 dihydrochloride | nAChR | Cholinergic receptor | Agonist | Probe | Ion channel |
| NCGC00017173-05 | Anabasine | nAChR | Cholinergic receptor | Agonist | Probe | Ion channel |
| NCGC00378583-01 | Ro 0437626 | P2X | Purinergic receptor type 2X | Antagonist | Probe | Ion channel |
| NCGC00387264-01 | JNJ 47965567 | P2X | Purinergic receptor type 2X | Antagonist | Probe | Ion channel |
| NCGC00370880-04 | A-740003 | P2X | Purinergic receptor type 2X | Antagonist | Probe | Ion channel |
| NCGC00347903-02 | Ro-51 | P2X | Purinergic receptor type 2X | Antagonist | Probe | Ion channel |
| NCGC00347904-03 | AZ-11645373 | P2X | Purinergic receptor type 2X | Antagonist | Probe | Ion channel |
| NCGC00484064-02 | AF-353 | P2X | Purinergic receptor type 2X | Antagonist | Probe | Ion channel |
| NCGC00347905-01 | AZ-10606120 | P2X | Purinergic receptor type 2X | Antagonist | Probe | Ion channel |
| NCGC00347909-03 | A-839977 | P2X | Purinergic receptor type 2X | Antagonist | Probe | Ion channel |
| NCGC00346721-03 | GW-791343 | P2X | Purinergic receptor type 2X | Antagonist | Probe | Ion channel |
| NCGC00370894-02 | 5-BDBD | P2X | Purinergic receptor type 2X | Antagonist | Probe | Ion channel |
| NCGC00685462-01 | Gefapixant | P2X | Purinergic receptor type 2X | Antagonist | Probe | Ion channel |
| NCGC00370698-02 | PSB-12062 | P2X | Purinergic receptor type 2X | Antagonist | Probe | Ion channel |
| NCGC00370701-03 | A-804598 | P2X | Purinergic receptor type 2X | Antagonist | Probe | Ion channel |
| NCGC00402268-01 | A-804598 | P2X | Purinergic receptor type 2X | Antagonist | Probe | Ion channel |
| NCGC00378772-01 | RO-3 | P2X | Purinergic receptor type 2X | Antagonist | Probe | Ion channel |
| NCGC00386667-01 | P2Y14 Antagonist Prodrug 7j hydrochloride | P2Y | P2Y receptor | Antagonist | Probe | Ion channel |
| NCGC00387209-01 | AR-C 118925XX | P2Y | P2Y receptor | Antagonist | Probe | Ion channel |
| NCGC00390624-03 | AZD-1283 | P2Y | P2Y receptor | Antagonist | Probe | Ion channel |
| NCGC00510466-01 | BPTU | P2Y | P2Y receptor | PAM | Probe | Ion channel |

| | | | | | | |
|---|---|---|---|---|---|---|
| NCGC00484059-02 | N6-(4-Hydroxybenzyl)adenosine | P2Y | P2Y receptor | Antagonist | Probe | Ion channel |
| NCGC00476206-01 | Yoda1 | PIEZO1 | Piezo type mechanosensitive ion channel component 1 | Agonist | Probe | Ion channel |
| NCGC00655664-01 | Dooku1 | PIEZO1 | Piezo type mechanosensitive ion channel component 1 | Antagonist | Probe | Ion channel |
| NCGC00383258-02 | Jedi2 | PIEZO1 | Piezo type mechanosensitive ion channel component 1 | Agonist | Probe | Ion channel |
| NCGC00382571-03 | Jedi1 | PIEZO1 | Piezo type mechanosensitive ion channel component 1 | Agonist | Probe | Ion channel |
| NCGC00370970-01 | Pyr3 | TRPA | TRP subfamily A member | Antagonist | Probe | Ion channel |
| NCGC00522005-01 | AM-0902 | TRPA | TRP subfamily A member | Antagonist | Probe | Ion channel |
| NCGC00378740-02 | HC-030031 | TRPA | TRP subfamily A member | Antagonist | Probe | Ion channel |
| NCGC00522548-01 | PF-4840154 | TRPC | TRP subfamily C member | Antagonist | Probe | Ion channel |
| NCGC00522452-01 | Pyr10 | TRPC | TRP subfamily C member | Antagonist | Probe | Ion channel |
| NCGC00390662-04 | Pyr6 | TRPC | TRP subfamily C member | Antagonist | Probe | Ion channel |
| NCGC00522486-01 | D-3263 | TRPC | TRP subfamily C member | Antagonist | Probe | Ion channel |
| NCGC00370959-02 | BCTC | TRPC | TRP subfamily C member | Antagonist | Probe | Ion channel |
| NCGC00285960-04 | Chembridge-5861528 | TRPC | TRP subfamily C member | Antagonist | Probe | Ion channel |
| NCGC00379231-02 | SN-2 | TRPC | TRP subfamily C member | Antagonist | Probe | Ion channel |
| NCGC00379188-04 | ML204 | TRPC | TRP subfamily C member | Antagonist | Probe | Ion channel |
| NCGC00685406-01 | HC-070 | TRPC4 | TRP subfamily C member 4 | Antagonist | Probe | Ion channel |
| NCGC00506826-01 | M-084 | TRPC4 | TRP subfamily C member 4 | Antagonist | Probe | Ion channel |
| NCGC00025073-06 | KB-R7943 | TRPC5 | TRP subfamily C member 5 | | Probe | Ion channel |
| NCGC00387486-02 | RQ-00203078 | TRPM | TRP subfamily M member | Antagonist | Probe | Ion channel |
| NCGC00387489-01 | TC-I 2014 | TRPM | TRP subfamily M member | Antagonist | Probe | Ion channel |
| NCGC00370691-04 | M8-B | TRPM | TRP subfamily M member | Antagonist | Probe | Ion channel |
| NCGC00379172-01 | TC-I 2000 | TRPM | TRP subfamily M member | Antagonist | Probe | Ion channel |
| NCGC00685397-01 | TRPM8 antagonist 2 | TRPM | TRP subfamily M member | Antagonist | Probe | Ion channel |
| NCGC00370824-02 | RN-1747 | TRPM | TRP subfamily M member | Antagonist | Probe | Ion channel |
| NCGC00387570-01 | 10-HDA | TRPM | TRP subfamily M member | Antagonist | Probe | Ion channel |
| NCGC00485432-01 | Umbellulone | TRPM | TRP subfamily M member | Antagonist | Probe | Ion channel |
| NCGC00480795-01 | Mifamurtide | TRPV | TRP subfamily V member | Agonist | Probe | Ion channel |
| NCGC00370767-03 | GSK-2193874 | TRPV | TRP subfamily V member | Antagonist | Probe | Ion channel |
| NCGC00250409-02 | GSK-1016790A | TRPV | TRP subfamily V member | Agonist | Probe | Ion channel |
| NCGC00263016-01 | DE-096 | TRPV | TRP subfamily V member | Antagonist | Probe | Ion channel |
| NCGC00685447-01 | Pico145 | TRPV | TRP subfamily V member | Antagonist | Probe | Ion channel |
| NCGC00263214-03 | HC-067047 | TRPV | TRP subfamily V member | Antagonist | Probe | Ion channel |
| NCGC00378585-02 | SAR-7334 | TRPV | TRP subfamily V member | Antagonist | Probe | Ion channel |
| NCGC00025031-06 | SKF-96365 | TRPV | TRP subfamily V member | Antagonist | Probe | Ion channel |
| NCGC00408813-01 | GSK-205 | TRPV | TRP subfamily V member | Antagonist | Probe | Ion channel |
| NCGC00015190-05 | Capsazepine | TRPV | TRP subfamily V member | Antagonist | Probe | Ion channel |
| NCGC00370826-01 | RN-1734 | TRPV | TRP subfamily V member | Antagonist | Probe | Ion channel |
| NCGC00388058-01 | N-(p-amylcinnamoyl) Anthranilic Acid | TRPV | TRP subfamily V member | Agonist | Probe | Ion channel |
| NCGC00386353-01 | Optovin | TRPV | TRP subfamily V member | Agonist | Probe | Ion channel |
| NCGC00025231-03 | SB-366791 | TRPV | TRP subfamily V member | Antagonist | Probe | Ion channel |
| NCGC00142360-05 | Podocarpic acid | TRPV | TRP subfamily V member | Agonist | Probe | Ion channel |
| NCGC00162411-05 | Arvanil | TRPV1 | TRP subfamily V member 1 | Agonist | Probe | Ion channel |

| | | | | | | |
|---|---|---|---|---|---|---|
| NCGC00161231-05 | NADA | TRPV1 | TRP subfamily V member 1 | Agonist | Probe | Ion channel |
| NCGC00685383-01 | A-1165442 | TRPV1 | TRP subfamily V member 1 | Antagonist | Probe | Ion channel |
| NCGC00402354-02 | JYL-1421 | TRPV1 | TRP subfamily V member 1 | Antagonist | Probe | Ion channel |
| NCGC00386568-01 | ABT-102 | TRPV1 | TRP subfamily V member 1 | Antagonist | Probe | Ion channel |
| NCGC00655634-01 | DPBA | TRPV1 | TRP subfamily V member 1 | Antagonist | Probe | Ion channel |
| NCGC00483055-01 | Astragaloside A | Cav | Voltage-gated calcium channel | Blocker | Probe | Ion channel |
| NCGC00162463-04 | Ionomycin | Cav | Voltage-gated calcium channel | activator | Probe | Ion channel |
| NCGC00025379-05 | SR-33805 | Cav | Voltage-gated calcium channel | Blocker | Probe | Ion channel |
| NCGC00346992-02 | Purfalcamine | Cav | Voltage-gated calcium channel | Blocker | Probe | Ion channel |
| NCGC00119853-04 | CDN1163 | Cav | Voltage-gated calcium channel | Blocker | Probe | Ion channel |
| NCGC00686689-01 | ACT-709478 | Cav | Voltage-gated calcium channel | Blocker | Probe | Ion channel |
| NCGC00344111-02 | YM-58483 | Cav | Voltage-gated calcium channel | Blocker | Probe | Ion channel |
| NCGC00389441-01 | NPS-2143 | Cav | Voltage-gated calcium channel | Blocker | Probe | Ion channel |
| NCGC00381746-03 | Calhex-231 | Cav | Voltage-gated calcium channel | Blocker | Probe | Ion channel |
| NCGC00686684-01 | TTA-Q6 | Cav | Voltage-gated calcium channel | Blocker | Probe | Ion channel |
| NCGC00402346-01 | GSK-7975A | Cav | Voltage-gated calcium channel | Blocker | Probe | Ion channel |
| NCGC00264075-01 | STS | Cav | Voltage-gated calcium channel | Blocker | Probe | Ion channel |
| NCGC00522540-03 | Calcium channel inhibitor 1 | Cav | Voltage-gated calcium channel | Blocker | Probe | Ion channel |
| NCGC00485404-01 | ONO-RS-082 | Cav | Voltage-gated calcium channel | Blocker | Probe | Ion channel |
| NCGC00686685-01 | Dehydronitrosonisoldipine | Cav | Voltage-gated calcium channel | Blocker | Probe | Ion channel |
| NCGC00378787-02 | Calcium-Sensing Receptor Antagonists I | Cav | Voltage-gated calcium channel | Blocker | Probe | Ion channel |
| NCGC00025210-07 | Bay K-8644 | Cav | Voltage-gated calcium channel | Blocker | Probe | Ion channel |
| NCGC00485485-01 | Ca2+ channel agonist 1 | Cav | Voltage-gated calcium channel | Blocker | Probe | Ion channel |
| NCGC00344505-01 | Calindol | Cav | Voltage-gated calcium channel | Blocker | Probe | Ion channel |
| NCGC00384735-01 | Columbianadin | Cav | Voltage-gated calcium channel | Blocker | Probe | Ion channel |
| NCGC00511374-02 | NS-638 | Cav | Voltage-gated calcium channel | Blocker | Probe | Ion channel |
| NCGC00380333-01 | Pimaric acid | Cav | Voltage-gated calcium channel | Blocker | Probe | Ion channel |
| NCGC00344509-02 | AC-265347 | Cav | Voltage-gated calcium channel | Blocker | Probe | Ion channel |
| NCGC00402329-02 | FKGK18 | Cav | Voltage-gated calcium channel | Blocker | Probe | Ion channel |
| NCGC00650290-01 | Methyl syringate | Cav | Voltage-gated calcium channel | Blocker | Probe | Ion channel |
| NCGC00163412-08 | Nigericin | Kv | Voltage-gated potassium channel | Blocker | Probe | Ion channel |
| NCGC00482986-01 | Daurisoline | Kv | Voltage-gated potassium channel | Blocker | Probe | Ion channel |
| NCGC00386732-01 | AVE-0118 | Kv | Voltage-gated potassium channel | Blocker | Probe | Ion channel |
| NCGC00390585-01 | Azimilide | Kv | Voltage-gated potassium channel | Blocker | Probe | Ion channel |
| NCGC00379200-05 | ML277 | Kv | Voltage-gated potassium channel | Activator | Probe | Ion channel |

| | | | | | | |
|---|---|---|---|---|---|---|
| NCGC00386772-01 | PK-THPP | Kv | Voltage-gated potassium channel | Blocker | Probe | Ion channel |
| NCGC00370962-01 | UK-78282 | Kv | Voltage-gated potassium channel | Blocker | Probe | Ion channel |
| NCGC00386602-01 | S-9947 | Kv | Voltage-gated potassium channel | Blocker | Probe | Ion channel |
| NCGC00387222-01 | HMR-1556 | Kv | Voltage-gated potassium channel | Blocker | Probe | Ion channel |
| NCGC00522484-01 | BMS-191095 | Kv | Voltage-gated potassium channel | Activator | Probe | Ion channel |
| NCGC00025301-03 | E-4031 | Kv | Voltage-gated potassium channel | Blocker | Probe | Ion channel |
| NCGC00378686-01 | L-364373 | Kv | Voltage-gated potassium channel | Activator | Probe | Ion channel |
| NCGC00522637-01 | GSK-369796 | Kv | Voltage-gated potassium channel | Blocker | Probe | Ion channel |
| NCGC00482765-01 | Rhynchophylline | Kv | Voltage-gated potassium channel | Blocker | Probe | Ion channel |
| NCGC00485529-01 | DFK | Kv | Voltage-gated potassium channel | Blocker | Probe | Ion channel |
| NCGC00371023-02 | VU-591 | Kv | Voltage-gated potassium channel | Blocker | Probe | Ion channel |
| NCGC00386909-04 | ML365 | Kv | Voltage-gated potassium channel | blocker | Probe | Ion channel |
| NCGC00017389-04 | Protopine | Kv | Voltage-gated potassium channel | Blocker | Probe | Ion channel |
| NCGC00165872-02 | PAP-1 | Kv | Voltage-gated potassium channel | Blocker | Probe | Ion channel |
| NCGC00015427-10 | FPL-64176 | Kv | Voltage-gated potassium channel | Activator | Probe | Ion channel |
| NCGC00165909-05 | TRAM-34 | Kv | Voltage-gated potassium channel | Blocker | Probe | Ion channel |
| NCGC00370935-03 | VU-0240551 | Kv | Voltage-gated potassium channel | Blocker | Probe | Ion channel |
| NCGC00379216-04 | NS-6180 | Kv | Voltage-gated potassium channel | Blocker | Probe | Ion channel |
| NCGC00390719-03 | 20-HETE | Kv | Voltage-gated potassium channel | Blocker | Probe | Ion channel |
| NCGC00379157-04 | ML133 | Kv | Voltage-gated potassium channel | Blocker | Probe | Ion channel |
| NCGC00402245-02 | HUP30 | Kv | Voltage-gated potassium channel | Activator | Probe | Ion channel |
| NCGC00686678-01 | ML402 | Kv | Voltage-gated potassium channel | Activator | Probe | Ion channel |
| NCGC00387162-01 | ICA-069673 | Kv | Voltage-gated potassium channel | Activator | Probe | Ion channel |
| NCGC00379147-04 | ML213 | Kv | Voltage-gated potassium channel | Activator | Probe | Ion channel |
| NCGC00686690-02 | GAL-021 | Kv | Voltage-gated potassium channel | Blocker | Probe | Ion channel |
| NCGC00018126-03 | Kavain | Kv | Voltage-gated potassium channel | Blocker | Probe | Ion channel |
| NCGC00686692-01 | SKA-121 | Kv | Voltage-gated potassium channel | Activator | Probe | Ion channel |
| NCGC00482999-01 | Bulleyaconicine A | Nav | Voltage-gated sodium channels | Blocker | Probe | Ion channel |
| NCGC00685381-01 | Sodium ionophore III | Nav | Voltage-gated sodium channels | Blocker | Probe | Ion channel |
| NCGC00370930-01 | A-887826 | Nav | Voltage-gated sodium channels | Blocker | Probe | Ion channel |
| NCGC00379156-01 | YM-244769 | Nav | Voltage-gated sodium channels | Blocker | Probe | Ion channel |
| NCGC00379083-02 | Sodium Channel inhibitor 1 | Nav | Voltage-gated sodium channels | Blocker | Probe | Ion channel |
| NCGC00510521-02 | AM-2099 | Nav | Voltage-gated sodium channels | Blocker | Probe | Ion channel |
| NCGC00370808-01 | KC-12291 | Nav | Voltage-gated sodium channels | Blocker | Probe | Ion channel |
| NCGC00379252-01 | ICA-121431 | Nav | Voltage-gated sodium channels | Blocker | Probe | Ion channel |

| | | | | | | |
|---|---|---|---|---|---|---|
| NCGC00686675-01 | BI-01383298 | Nav | Voltage-gated sodium channels | Blocker | Probe | Ion channel |
| NCGC00168796-02 | Rhodotoxin | Nav | Voltage-gated sodium channels | Blocker | Probe | Ion channel |
| NCGC00484992-01 | Veratramine | Nav | Voltage-gated sodium channels | Blocker | Probe | Ion channel |
| NCGC00685456-01 | PF-06869206 | Nav | Voltage-gated sodium channels | Blocker | Probe | Ion channel |
| NCGC00378867-04 | SEA0400 | Nav | Voltage-gated sodium channels | Blocker | Probe | Ion channel |
| NCGC00015147-11 | Benzamil | Nav | Voltage-gated sodium channels | Blocker | Probe | Ion channel |
| NCGC00485395-01 | ORM-10103 | Nav | Voltage-gated sodium channels | Blocker | Probe | Ion channel |
| NCGC00408812-05 | GS-967 | Nav | Voltage-gated sodium channels | Blocker | Probe | Ion channel |
| NCGC00015108-12 | HMA-5 | Nav | Voltage-gated sodium channels | Blocker | Probe | Ion channel |
| NCGC00387094-04 | EIPA | Nav | Voltage-gated sodium channels | Blocker | Probe | Ion channel |
| NCGC00379232-02 | PF-04885614 | Nav | Voltage-gated sodium channels | Blocker | Probe | Ion channel |
| NCGC00390339-01 | Veratridine | Nav1.7 | Sodium voltage-gated channel alpha subunit 9 | Blocker | Probe | Ion channel |
| NCGC00379126-01 | Nav1.7 blocker 52 | Nav1.7 | Sodium voltage-gated channel alpha subunit 9 | Blocker | Probe | Ion channel |
| NCGC00386641-01 | Nav1.7 blocker 24 | Nav1.7 | Sodium voltage-gated channel alpha subunit 9 | Blocker | Probe | Ion channel |
| NCGC00685382-01 | GNE-131 | Nav1.7 | Sodium voltage-gated channel alpha subunit 9 | Blocker | Probe | Ion channel |
| NCGC00387182-01 | Nav 26 | Nav1.7 | Sodium voltage-gated channel alpha subunit 9 | Blocker | Probe | Ion channel |
| NCGC00378627-02 | Nav1.7 inhibitor | Nav1.7 | Sodium voltage-gated channel alpha subunit 9 | Blocker | Probe | Ion channel |
| NCGC00685357-01 | Nav1.7-IN-2 | Nav1.7 | Sodium voltage-gated channel alpha subunit 9 | Blocker | Probe | Ion channel |
| NCGC00386684-01 | XEN-907 | Nav1.7 | Sodium voltage-gated channel alpha subunit 9 | Blocker | Probe | Ion channel |
| NCGC00655639-01 | N-Me-aminopyrimidinone 9 | Nav1.7 | Sodium voltage-gated channel alpha subunit 9 | Blocker | Probe | Ion channel |
| NCGC00522583-01 | PF-01247324 | Nav1.8 | Sodium voltage-gated channel alpha subunit 10 | Blocker | Probe | Ion channel |
| NCGC00685413-01 | AZD-7594 | GCR | Glucocorticoid receptor | Agonist | Probe | Nuclear receptor |
| NCGC00685350-01 | AZD-9567 | GCR | Glucocorticoid receptor | Agonist | Probe | Nuclear receptor |
| NCGC00482877-01 | g-PPT | GCR | Glucocorticoid receptor | Agonist | Probe | Nuclear receptor |
| NCGC00685405-01 | Desisobutyryl-ciclesonide | GCR | Glucocorticoid receptor | Agonist | Probe | Nuclear receptor |
| NCGC00685369-01 | AZD-2906 | GCR | Glucocorticoid receptor | Agonist | Probe | Nuclear receptor |
| NCGC00358128-03 | AL-082D06 | GCR | Glucocorticoid receptor | Antagonist | Probe | Nuclear receptor |
| NCGC00386284-02 | AZD-3514 | NR | Nuclear Receptor | Antagonist | Probe | Nuclear receptor |
| NCGC00482601-01 | Patchouli | NR | Nuclear Receptor | Agonist | Probe | Nuclear receptor |
| NCGC00483005-01 | 20(S)-Ginsenoside Rh1 | PPAR | Peroxisome proliferator-activated receptor | Agonist | Probe | Nuclear receptor |
| NCGC00685400-01 | NXT-629 | PPAR | Peroxisome proliferator-activated receptor | Antagonist | Probe | Nuclear receptor |
| NCGC00095894-04 | Astaxanthin | PPAR | Peroxisome proliferator-activated receptor | Agonist | Probe | Nuclear receptor |
| NCGC00387450-01 | CDDO-Im | PPAR | Peroxisome proliferator-activated receptor | Agonist | Probe | Nuclear receptor |

| | | | | | | |
|---|---|---|---|---|---|---|
| NCGC00025251-09 | GW-1929 | PPAR | Peroxisome proliferator-activated receptor | Agonist | Probe | Nuclear receptor |
| NCGC00588880-01 | Lanifibranor | PPAR | Peroxisome proliferator-activated receptor | Agonist | Probe | Nuclear receptor |
| NCGC00496834-01 | AZ-6102 | PPAR | Peroxisome proliferator-activated receptor | Agonist | Probe | Nuclear receptor |
| NCGC00476202-01 | DG172 | PPAR | Peroxisome proliferator-activated receptor | Antagonist | Probe | Nuclear receptor |
| NCGC00370933-07 | FH-535 | PPAR | Peroxisome proliferator-activated receptor | Antagonist | Probe | Nuclear receptor |
| NCGC00163517-02 | Bavachinin | PPAR | Peroxisome proliferator-activated receptor | Agonist | Probe | Nuclear receptor |
| NCGC00650293-01 | Phytol | PPAR | Peroxisome proliferator-activated receptor | Agonist | Probe | Nuclear receptor |
| NCGC00161609-04 | Magnolol | PPAR | Peroxisome proliferator-activated receptor | Agonist | Probe | Nuclear receptor |
| NCGC00482508-01 | Gypenoside XLIX | PPARa | Peroxisome proliferator-activated receptor alpha | Agonist | Probe | Nuclear receptor |
| NCGC00482527-01 | Anemoside B4 | ILR | Interleukin receptor | Antagonist | Probe | Catallytic receptor |
| NCGC00522470-01 | OSU-T315 | ILR | Interleukin receptor | Antagonist | Probe | Catallytic receptor |
| NCGC00248064-06 | CP-456773 | ILR | Interleukin receptor | Antagonist | Probe | Catallytic receptor |
| NCGC00356071-15 | CPI-203 | ILR | Interleukin receptor | Antagonist | Probe | Catallytic receptor |
| NCGC00025256-02 | YM-90709 | ILR | Interleukin receptor | Antagonist | Probe | Catallytic receptor |
| NCGC00590984-01 | CA-4948 | TLR | Toll-like receptor | Antagonist | Probe | Catallytic receptor |
| NCGC00387208-03 | CU-CPT-22 | TLR | Toll-like receptor | Agonist | Probe | Catallytic receptor |
| NCGC00482674-01 | Leonurine | TLR | Toll-like receptor | Antagonist | Probe | Catallytic receptor |
| NCGC00387491-01 | CU-T12-9 | TLR2 | Toll-like receptor 2 | Agonist | Probe | Catallytic receptor |
| NCGC00685339-01 | C-29 | TLR2 | Toll-like receptor 2 | Antagonist | Probe | Catallytic receptor |
| NCGC00685420-01 | CU-CPT-17e | TLR3 | Toll-like receptor 3 | Agonist | Probe | Catallytic receptor |
| NCGC00379226-01 | CU-CPT-4a | TLR3 | Toll-like receptor 3 | Antagonist | Probe | Catallytic receptor |
| NCGC00685379-01 | Telratolimod | TLR7 | Toll-like receptor 7 | Antagonist | Probe | Catallytic receptor |
| NCGC00387452-01 | DSR-6434 | TLR7 | Toll-like receptor 7 | Agonist | Probe | Catallytic receptor |
| NCGC00685437-01 | HY-103698A | TLR7 | Toll-like receptor 7 | Agonist | Probe | Catallytic receptor |
| NCGC00685399-01 | HY-103039 | TLR7 | Toll-like receptor 7 | Agonist | Probe | Catallytic receptor |
| NCGC00387769-01 | Gardiquimod | TLR7 | Toll-like receptor 7 | Agonist | Probe | Catallytic receptor |
| NCGC00378688-02 | W-5494 | TLR8 | Toll-like receptor 8 | Agonist | Probe | Catallytic receptor |
| NCGC00685431-01 | CU-CPT-9a | TLR8 | Toll-like receptor 8 | Antagonist | Probe | Catallytic receptor |
| NCGC00685348-01 | CU-CPT-8m | TLR8 | Toll-like receptor 8 | Antagonist | Probe | Catallytic receptor |
| NCGC00685370-01 | CU-CPT-9b | TLR8 | Toll-like receptor 8 | Antagonist | Probe | Catallytic receptor |
| NCGC00485947-01 | E6446 | TLR9 | Toll-like receptor 8 | Antagonist | Probe | Catallytic receptor |
| NCGC00482526-01 | Platycodin D | TNFa | Tumor necrosis factor | Antagonist | Probe | Catallytic receptor |
| NCGC00482991-01 | Hypaconitine | TNFa | Tumor necrosis factor | Antagonist | Probe | Catallytic receptor |
| NCGC00476197-01 | C-87 | TNFa | Tumor necrosis factor | Antagonist | Probe | Catallytic receptor |

| | | | | | | |
|---|---|---|---|---|---|---|
| NCGC00169046-04 | Prim-o-glucosylcimifugin | TNFa | Tumor necrosis factor | Antagonist | Probe | Catallytic receptor |
| NCGC00096001-03 | fMLP | TNFa | Tumor necrosis factor | Antagonist | Probe | Catallytic receptor |
| NCGC00482823-02 | Peimine | TNFa | Tumor necrosis factor | Antagonist | Probe | Catallytic receptor |
| NCGC00387493-02 | R-7050 | TNFa | Tumor necrosis factor | Antagonist | Probe | Catallytic receptor |
| NCGC00685443-01 | Hispidol | TNFa | Tumor necrosis factor | Antagonist | Probe | Catallytic receptor |
| NCGC00685466-01 | DCVC | TNFa | Tumor necrosis factor | Antagonist | Probe | Catallytic receptor |
| NCGC00485142-01 | Picrotoxinin | GlyT | Glycine transporter | Inhibitor | Probe | Transporter |
| NCGC00247954-03 | SSR-504734 | GlyT1 | Glycine transporter 1 | Inhibitor | Probe | Transporter |
| NCGC00370805-01 | LY-2365109 | GlyT1 | Glycine transporter 1 | Inhibitor | Probe | Transporter |
| NCGC00387477-01 | ASP-2535 | GlyT1 | Glycine transporter 1 | Inhibitor | Probe | Transporter |
| NCGC00379203-01 | ORG-25543 | GlyT2 | Glycine transporter 2 | Inhibitor | Probe | Transporter |
| NCGC00650299-01 | Ethyl (triphenylphosphoranylidene) acetate | AChE | Acetylcholinesterase | Inhibitor | Probe | Enzyme |
| NCGC00247622-02 | Jatrorrhizine | AChE | Acetylcholinesterase | Inhibitor | Probe | Enzyme |
| NCGC00163413-07 | Lycorine | AChE | Acetylcholinesterase | Inhibitor | Probe | Enzyme |
| NCGC00482662-01 | Dehydroevodiamine | AChE | Acetylcholinesterase | Inhibitor | Probe | Enzyme |
| NCGC00016080-05 | Vesamicol | AChE | Acetylcholinesterase | Inhibitor | Probe | Enzyme |
| NCGC00017221-13 | Harmaline | AChE | Acetylcholinesterase | Inhibitor | Probe | Enzyme |
| NCGC00183866-01 | Oxypertine | AC1 | Adenylate Cyclase 1 | Inhibitor | Probe | Enzyme |
| NCGC00522009-01 | ST-034307 | AC1 | Adenylate Cyclase 1 | Inhibitor | Probe | Enzyme |
| NCGC00685427-01 | CB-7921220 | AC1 | Adenylate Cyclase 1 | Inhibitor | Probe | Enzyme |
| NCGC00015912-08 | SQ22536 | AC1 | Adenylate Cyclase 1 | Inhibitor | Probe | Enzyme |
| NCGC00480789-01 | Ginsenoside C-K | COX | Cyclooxygenase | Inhibitor | Probe | Enzyme |
| NCGC00482874-01 | Echinocystic acid | COX | Cyclooxygenase | Inhibitor | Probe | Enzyme |
| NCGC00169402-02 | Isoorientin | COX | Cyclooxygenase | Inhibitor | Probe | Enzyme |
| NCGC00482733-01 | Columbin | COX | Cyclooxygenase | Inhibitor | Probe | Enzyme |
| NCGC00015933-05 | SC-560 | COX | Cyclooxygenase | Inhibitor | Probe | Enzyme |
| NCGC00163514-01 | Aucubin | COX | Cyclooxygenase | Inhibitor | Probe | Enzyme |
| NCGC00179896-02 | Coniferin | COX | Cyclooxygenase | Inhibitor | Probe | Enzyme |
| NCGC00378722-05 | FK-3311 | COX | Cyclooxygenase | Inhibitor | Probe | Enzyme |
| NCGC00482701-01 | Jaceosidin | COX | Cyclooxygenase | Inhibitor | Probe | Enzyme |
| NCGC00181008-03 | Olsalazine sodium | COX | Cyclooxygenase | Inhibitor | Probe | Enzyme |
| NCGC00385881-02 | Inulicin | COX | Cyclooxygenase | Inhibitor | Probe | Enzyme |
| NCGC00685347-01 | Hexahydrocurcumin | COX | Cyclooxygenase | Inhibitor | Probe | Enzyme |
| NCGC00482571-01 | Syringaldehyde | COX | Cyclooxygenase | Inhibitor | Probe | Enzyme |
| NCGC00174656-02 | Phenidone | COX | Cyclooxygenase | Inhibitor | Probe | Enzyme |
| NCGC00025192-04 | FR122047 | COX1 | Cyclooxygenase 1 | Inhibitor | Probe | Enzyme |
| NCGC00655662-01 | 2-[1-(4-chlorobenzoyl)-5-methoxy-2-methyl-1H-indol-3-yl]-n-[(1R)-1-(hydroxymethyl)propyl]acetamide | COX1 | Cyclooxygenase 1 | Inhibitor | Probe | Enzyme |
| NCGC00370982-01 | SC-236 | COX1 | Cyclooxygenase 1 | Inhibitor | Probe | Enzyme |
| NCGC00655646-01 | FK-881 | COX1 | Cyclooxygenase 1 | Inhibitor | Probe | Enzyme |
| NCGC00095907-06 | Nonoxynol-9 | COX2 | Cyclooxygenase 2 | Inhibitor | Probe | Enzyme |

| | | | | | | |
|---|---|---|---|---|---|---|
| NCGC00531680-02 | Beta-Elemonic | COX2 | Cyclooxygenase 2 | Inhibitor | Probe | Enzyme |
| NCGC00168853-11 | Lupeol | COX2 | Cyclooxygenase 2 | Inhibitor | Probe | Enzyme |
| NCGC00015900-02 | Seliciclib | COX2 | Cyclooxygenase 2 | Inhibitor | Probe | Enzyme |
| NCGC00355143-02 | Dehydrodiisoeugenol | COX2 | Cyclooxygenase 2 | Inhibitor | Probe | Enzyme |
| NCGC00169750-03 | 8-Gingerol | COX2 | Cyclooxygenase 2 | Inhibitor | Probe | Enzyme |
| NCGC00181152-02 | Cisplatin | COX2 | Cyclooxygenase 2 | Inhibitor | Probe | Enzyme |
| NCGC00485328-01 | Sapropterin | COX2 | Cyclooxygenase 2 | Inhibitor | Probe | Enzyme |
| NCGC00160337-02 | Tryptanthrin | COX2 | Cyclooxygenase 2 | Inhibitor | Probe | Enzyme |
| NCGC00482598-01 | alpha-Cyperone | COX2 | Cyclooxygenase 2 | Inhibitor | Probe | Enzyme |
| NCGC00482587-01 | Ethyl caffeate | COX2 | Cyclooxygenase 2 | Inhibitor | Probe | Enzyme |
| NCGC00482584-01 | Ethyl 4-Methoxycinnamate | COX2 | Cyclooxygenase 2 | Inhibitor | Probe | Enzyme |
| NCGC00346837-05 | JZL-195 | FAAH | Fatty acid amide hydrolase | Inhibitor | Probe | Enzyme |
| NCGC00522488-01 | JNJ-42165279 | FAAH | Fatty acid amide hydrolase | Inhibitor | Probe | Enzyme |
| NCGC00485886-03 | FAAH-IN-2 | FAAH | Fatty acid amide hydrolase | Inhibitor | Probe | Enzyme |
| NCGC00159571-03 | LY-2183240 | FAAH | Fatty acid amide hydrolase | Inhibitor | Probe | Enzyme |
| NCGC00378930-01 | TUG-770 | FAAH | Fatty acid amide hydrolase | Agonist | Probe | Enzyme |
| NCGC00505864-01 | BIA 10-2474 | FAAH | Fatty acid amide hydrolase | Inhibitor | Probe | Enzyme |
| NCGC00167723-05 | 2-PMPA | GCPII | Folate hydrolase prostate-specific membrane antigen 1 | Inhibitor | Probe | Enzyme |
| NCGC00688028-01 | 2-MPPA | GCPII | Folate hydrolase prostate-specific membrane antigen 1 | Inhibitor | Probe | Enzyme |
| NCGC00024489-06 | Quisqualate | GCPII | Folate hydrolase prostate-specific membrane antigen 1 | Inhibitor | Probe | Enzyme |
| NCGC00346840-03 | Tubacin | HDAC | Histone deacetylase | Inhibitor | Probe | Enzyme |
| NCGC00510505-02 | WT-161 | HDAC | Histone deacetylase | Inhibitor | Probe | Enzyme |
| NCGC00250390-03 | CAY10603 | HDAC | Histone deacetylase | Inhibitor | Probe | Enzyme |
| NCGC00510486-02 | EDO-S101 | HDAC | Histone deacetylase | Inhibitor | Probe | Enzyme |
| NCGC00522526-01 | HDAC-IN-4 | HDAC | Histone deacetylase | Inhibitor | Probe | Enzyme |
| NCGC00522525-01 | HDAC-IN-3 | HDAC | Histone deacetylase | Inhibitor | Probe | Enzyme |
| NCGC00685385-01 | HDAC6-IN-1 | HDAC | Histone deacetylase | Inhibitor | Probe | Enzyme |
| NCGC00345786-02 | HDAC-IN-4 | HDAC | Histone deacetylase | Inhibitor | Probe | Enzyme |
| NCGC00165855-06 | Oxamflatin | HDAC | Histone deacetylase | Inhibitor | Probe | Enzyme |
| NCGC00263618-04 | BML-210 | HDAC | Histone deacetylase | Inhibitor | Probe | Enzyme |
| NCGC00263602-04 | TC-H 106 | HDAC | Histone deacetylase | Inhibitor | Probe | Enzyme |
| NCGC00531749-01 | ACY-775 | HDAC | Histone deacetylase | Inhibitor | Probe | Enzyme |
| NCGC00522469-01 | NKL-22 | HDAC | Histone deacetylase | Inhibitor | Probe | Enzyme |
| NCGC00685336-01 | SR-4370 | HDAC | Histone deacetylase | Inhibitor | Probe | Enzyme |
| NCGC00408830-02 | BRD-73954 | HDAC | Histone deacetylase | Inhibitor | Probe | Enzyme |
| NCGC00511370-02 | ACY-738 | HDAC | Histone deacetylase | Inhibitor | Probe | Enzyme |
| NCGC00689778-01 | Vafidemstat | MAO | Monoamine oxidase | inhibitor | Probe | Enzyme |
| NCGC00385504-01 | Rosiridin | MAO | Monoamine oxidase | Inhibitor | Probe | Enzyme |
| NCGC00485217-01 | 8-CSC | MAO | Monoamine oxidase | Inhibitor | Probe | Enzyme |
| NCGC00370705-03 | PF9601N | MAO | Monoamine oxidase | Inhibitor | Probe | Enzyme |
| NCGC00017269-09 | Formononetin | MAO | Monoamine oxidase | Inhibitor | Probe | Enzyme |
| NCGC00485141-01 | M-Mptp | MAO | Monoamine oxidase | Inhibitor | Probe | Enzyme |
| NCGC00246983-02 | Isatin | MAO | Monoamine oxidase | Inhibitor | Probe | Enzyme |
| NCGC00371031-01 | AR-C 102222 | NOS | Nitric oxide synthase | Inhibitor | Probe | Enzyme |
| NCGC00370813-03 | ARL-17477 | NOS | Nitric oxide synthase | Inhibitor | Probe | Enzyme |

| | | | | | | |
|---|---|---|---|---|---|---|
| NCGC00487203-01 | ZL-006 | NOS | Nitric oxide synthase | Inhibitor | Probe | Enzyme |
| NCGC00522450-01 | IC-87201 | NOS | Nitric oxide synthase | Inhibitor | Probe | Enzyme |
| NCGC00378738-01 | BYK-191023 | NOS | Nitric oxide synthase | Inhibitor | Probe | Enzyme |
| NCGC00476011-01 | 6-Biopterin | NOS | Nitric oxide synthase | Inhibitor | Probe | Enzyme |
| NCGC00016095-10 | 1400W | NOS | Nitric oxide synthase | Inhibitor | Probe | Enzyme |
| NCGC00015664-06 | (S)-Methylisothiourea | NOS | Nitric oxide synthase | Inhibitor | Probe | Enzyme |
| NCGC00485944-02 | TAPI-1 | ADAM17/TACE | ADAM17/TACE | Inhibitor | Probe | Enzyme |
| NCGC00015953-06 | SCH-28080 | ATPase | ATPase | Inhibitor | Probe | Enzyme |
| NCGC00025177-18 | Suramin | GCPII | GCPII | Inhibitor | Probe | Enzyme |
| NCGC00378687-03 | Verbascoside | NOS | Nitric oxide synthase | Inhibitor | Probe | Enzyme |
| NCGC00482810-01 | BMS-202 | PD-L1 | PD-L1 | Inhibitor | Probe | Enzyme |
| NCGC00015141-07 | Bromoenol | PLA2 | phospholipase A4 | Inhibitor | Probe | Enzyme |
| NCGC00015672-17 | Morin | PTP1B | PTP1B | Inhibitor | Probe | Enzyme |
| NCGC00390560-02 | IT-I214 | PDE | cAMP-specific 3';5'-cyclic phosphodiesterase | inhibitor | Probe | Enzyme |
| NCGC00532447-01 | Ziritaxestat | PDE | cAMP-specific 3';5'-cyclic phosphodiesterase | inhibitor | Probe | Enzyme |
| NCGC00522449-01 | Autotaxin modulator 1 | PDE | cAMP-specific 3';5'-cyclic phosphodiesterase | inhibitor | Probe | Enzyme |
| NCGC00387145-01 | HA130 | PDE | cAMP-specific 3';5'-cyclic phosphodiesterase | inhibitor | Probe | Enzyme |
| NCGC00685361-01 | TP-10 | PDE | cAMP-specific 3';5'-cyclic phosphodiesterase | inhibitor | Probe | Enzyme |
| NCGC00248048-02 | CP-671305 | PDE | cAMP-specific 3';5'-cyclic phosphodiesterase | inhibitor | Probe | Enzyme |
| NCGC00685398-01 | PF-05180999 | PDE | cAMP-specific 3';5'-cyclic phosphodiesterase | inhibitor | Probe | Enzyme |
| NCGC00165289-05 | ML030 | PDE | cAMP-specific 3';5'-cyclic phosphodiesterase | inhibitor | Probe | Enzyme |
| NCGC00522529-01 | PF-04957325 | PDE | cAMP-specific 3';5'-cyclic phosphodiesterase | inhibitor | Probe | Enzyme |
| NCGC00248057-02 | CI-1044 | PDE | cAMP-specific 3';5'-cyclic phosphodiesterase | inhibitor | Probe | Enzyme |
| NCGC00685417-01 | PDE1-IN-2 | PDE | cAMP-specific 3';5'-cyclic phosphodiesterase | inhibitor | Probe | Enzyme |
| NCGC00379038-01 | Nortadalafil | PDE | cAMP-specific 3';5'-cyclic phosphodiesterase | inhibitor | Probe | Enzyme |
| NCGC00510194-01 | PDE10-IN-1 | PDE | cAMP-specific 3';5'-cyclic phosphodiesterase | inhibitor | Probe | Enzyme |
| NCGC00165747-03 | BAY 73-6691 | PDE | cAMP-specific 3';5'-cyclic phosphodiesterase | inhibitor | Probe | Enzyme |
| NCGC00024587-06 | MY-5445 | PDE | cAMP-specific 3';5'-cyclic phosphodiesterase | inhibitor | Probe | Enzyme |
| NCGC00685389-01 | BI-409306 | PDE | cAMP-specific 3';5'-cyclic phosphodiesterase | inhibitor | Probe | Enzyme |
| NCGC00685421-01 | BW-A 78U | PDE | cAMP-specific 3';5'-cyclic phosphodiesterase | inhibitor | Probe | Enzyme |
| NCGC00168459-04 | NCGC00168459 | PDE4A | Phosphodiesterase 4A | Inhibitor | Probe | Enzyme |
| NCGC00408889-01 | Eggmanone | PDE4A | Phosphodiesterase 4A | Inhibitor | Probe | Enzyme |
| NCGC00370829-01 | RS-25344 | PDE4A | Phosphodiesterase 4A | Inhibitor | Probe | Enzyme |
| NCGC00387127-01 | CDP840 | PDE4A | Phosphodiesterase 4A | Inhibitor | Probe | Enzyme |
| NCGC00250394-01 | NVP-ABE171 | PDE4A | Phosphodiesterase 4A | Inhibitor | Probe | Enzyme |
| NCGC00025308-03 | YM-976 | PDE4A | Phosphodiesterase 4A | Inhibitor | Probe | Enzyme |
| NCGC00015166-11 | Ro 20-1724 | PDE4A | Phosphodiesterase 4A | Inhibitor | Probe | Enzyme |
| NCGC00371101-01 | MRK-560 | G-secretase | Gamma-secretase | Inhibitor | Probe | Enzyme |

| | | | | | | |
|---|---|---|---|---|---|---|
| NCGC00378720-02 | GSM1 | G-secretase | Gamma-secretase | Inhibitor | Probe | Enzyme |
| NCGC00346875-02 | BMS-299897 | G-secretase | Gamma-secretase | Inhibitor | Probe | Enzyme |
| NCGC00386563-01 | gamma-Secretase Inhibitor BZ | G-secretase | Gamma-secretase | Inhibitor | Probe | Enzyme |
| NCGC00485942-01 | QCR-41 | G-secretase | Gamma-secretase | Inhibitor | Probe | Enzyme |
| NCGC00370685-03 | JNJ-40418677 | G-secretase | Gamma-secretase | Inhibitor | Probe | Enzyme |
| NCGC00378929-01 | CS-1149 | G-secretase | Gamma-secretase | Inhibitor | Probe | Enzyme |
| NCGC00378817-02 | E-2012 | G-secretase | Gamma-secretase | Inhibitor | Probe | Enzyme |
| NCGC00378743-01 | CS-0329 | G-secretase | Gamma-secretase | Inhibitor | Probe | Enzyme |
| NCGC00346883-05 | BIO-acetoxime | GSK-3 | Glycogen synthase kinase-3 beta | Inhibitor | Probe | Kinase |
| NCGC00015950-12 | SB-415286 | GSK-3 | Glycogen synthase kinase-3 beta | Inhibitor | Probe | Kinase |
| NCGC00482731-01 | BIO | GSK-3 | Glycogen synthase kinase-3 beta | Inhibitor | Probe | Kinase |
| NCGC00346638-04 | TWS-119 | GSK-3 | Glycogen synthase kinase-3 beta | Inhibitor | Probe | Kinase |
| NCGC00522478-01 | CCG-215022 | GRK | GRK | Inhibitor | Probe | Kinase |
| NCGC00379050-01 | IRAK inhibitor 6 | IRAK | IRAK | Inhibitor | Probe | Kinase |
| NCGC00186035-06 | IRAK-1-4 Inhibitor I | IRAK | IRAK | Inhibitor | Probe | Kinase |
| NCGC00601693-01 | Takinib | TAK1 | TAK1 | Inhibitor | Probe | Kinase |
| NCGC00242484-04 | WYE-354 | mTOR | Mechanistic target of rapamycin kinase | Inhibitor | Probe | Kinase |
| NCGC00507877-02 | CZ415 | mTOR | Mechanistic target of rapamycin kinase | Inhibitor | Probe | Kinase |
| NCGC00685414-01 | PQR620 | mTOR | Mechanistic target of rapamycin kinase | Inhibitor | Probe | Kinase |
| NCGC00370729-05 | MHY1485 | mTOR | Mechanistic target of rapamycin kinase | Inhibitor | Probe | Kinase |
| NCGC00346635-02 | WYE-125132 | mTORC1 | Mechanistic target of rapamycin kinase | Inhibitor | Probe | Kinase |
| NCGC00250386-09 | BX-795 | IKK | IKB kinase | Inhibitor | Probe | Kinase |
| NCGC00481567-02 | LY-2409881 | IKK | IKB kinase | Inhibitor | Probe | Kinase |
| NCGC00486904-01 | MRT-68601 | IKK | IKB kinase | Inhibitor | Probe | Kinase |
| NCGC00242056-04 | IKK-3 Inhibitor IX | IKK | IKB kinase | Inhibitor | Probe | Kinase |
| NCGC00481403-01 | AZD-3264 | IKK | IKB kinase | Inhibitor | Probe | Kinase |
| NCGC00506805-01 | BI-605906 | IKK | IKB kinase | Inhibitor | Probe | Kinase |
| NCGC00522045-02 | IKK-IN-1 | IKK | IKB kinase | Inhibitor | Probe | Kinase |
| NCGC00347659-02 | Tenuifoliside A | NFKB | Nuclear factor kappa-light-chain-enhancer of activated B cells | Inhibitor | Probe | Kinase |
| NCGC00250384-02 | PHA-408 | NFKB | Nuclear factor kappa-light-chain-enhancer of activated B cells | Inhibitor | Probe | Kinase |
| NCGC00390255-03 | Maslinic acid | NFKB | Nuclear factor kappa-light-chain-enhancer of activated B cells | Inhibitor | Probe | Kinase |
| NCGC00180826-02 | Praeruptorin A | NFKB | Nuclear factor kappa-light-chain-enhancer of activated B cells | Inhibitor | Probe | Kinase |
| NCGC00344390-04 | QNZ | NFKB | Nuclear factor kappa-light-chain-enhancer of activated B cells | Inhibitor | Probe | Kinase |
| NCGC00163567-08 | Honokiol | NFKB | Nuclear factor kappa-light-chain-enhancer of activated B cells | Inhibitor | Probe | Kinase |
| NCGC00537961-01 | Curcumenol | NFKB | Nuclear factor kappa-light-chain-enhancer of activated B cells | Inhibitor | Probe | Kinase |
| NCGC00178332-02 | Genkwanin | MAPK | Mitogen-activated protein kinase | Inhibitor | Probe | Kinase |

| | | | | | | |
|---|---|---|---|---|---|---|
| NCGC00601699-01 | Lazertinib | EGFR | Epidermal growth factor receptor | Inhibitor | Probe | Kinase |
| NCGC00483922-02 | AZD-3759 | EGFR | Epidermal growth factor receptor | Inhibitor | Probe | Kinase |
| NCGC00651749-01 | Theliatinib | EGFR | Epidermal growth factor receptor | Inhibitor | Probe | Kinase |
| NCGC00685416-01 | CH7057288 | TrkA | Tropomyosin receptor kinase | Inhibitor | Probe | Kinase |
| NCGC00346936-03 | GNF-5837 | TrkA | Tropomyosin receptor kinase | Inhibitor | Probe | Kinase |
| NCGC00507859-03 | Larotrectinib | TrkA | Tropomyosin receptor kinase | Inhibitor | Probe | Kinase |
| NCGC00250381-03 | AZ-23 | TrkA | Tropomyosin receptor kinase | Inhibitor | Probe | Kinase |
| NCGC00601829-01 | Selitrectinib | TrkA | Tropomyosin receptor kinase | Inhibitor | Probe | Kinase |
| NCGC00242051-03 | GW-2580 | TrkA | Tropomyosin receptor kinase | Inhibitor | Probe | Kinase |
| NCGC00510692-02 | Repotrectinib | TrkA | Tropomyosin receptor kinase | Inhibitor | Probe | Kinase |
| NCGC00499184-01 | LM22B-10 | TrkB | Tropomyosin receptor kinase | Inhibitor | Probe | Kinase |
| NCGC00379202-02 | ANA-12 | TrkB | Tropomyosin receptor kinase | Inhibitor | Probe | Kinase |
| NCGC00387179-02 | LM22A-4 | TrkB | Tropomyosin receptor kinase | Inhibitor | Probe | Kinase |
| NCGC00095217-05 | Tropoflavin | TrkB | Tropomyosin receptor kinase | activator | Probe | Kinase |
| NCGC00388216-01 | Corilagin | carbonic anhydrase | carbonic anhydrase | Inhibitor | Probe | Others |
| NCGC00483017-01 | Thymopentin | cytokines | cytokines | activator | Probe | Others |
| NCGC00387254-04 | BMS-309403 | FABP4 | fatty acid binding protein 4 | Inhibitor | Probe | Others |
| NCGC00508899-01 | MCC-950 | NLRP3 | NLRP3 | inhibitor | Probe | Others |
| NCGC00686539-01 | CY-09 | NLRP3 | NLRP3 | inhibitor | Probe | Others |
| NCGC00482961-01 | Dehydroandrographolide | TMEM16A | TMEM16A | Inhibitor | Probe | Others |
